# Regulation of DNA damage response by trimeric G-protein Signaling

**DOI:** 10.1101/2021.07.21.452842

**Authors:** Amer Ali Abd El-Hafeez, Nina Sun, Anirban Chakraborty, Jason Ear, Suchismita Roy, Pranavi Chamarthi, Navin Rajapakse, Soumita Das, Kathryn E. Luker, Tapas K. Hazra, Gary D. Luker, Pradipta Ghosh

## Abstract

Upon sensing DNA double-strand breaks (DSBs), eukaryotic cells either die or repair DSBs *via* one of two competing pathways, i.e., non-homologous end-joining (NHEJ) or homologous recombination (HR). We show that cell fate after DNA damage hinges on the guanine nucleotide-exchange modulator of heterotrimeric G-protein, Giα•βγ, GIV/Girdin. GIV suppresses HR by binding and sequestering BRCA1, a key coordinator of multiple steps within the HR pathway, away from DSBs; it does so using a C-terminal motif that binds BRCA1’s BRCT-modules *via* both phospho-dependent and -independent mechanisms. GIV promotes NHEJ, and binds and activates Gi and enhances the ‘free’ Gβγ→PI-3-kinase→Akt pathway, thus revealing the enigmatic origin of prosurvival Akt signals during dsDNA repair. Absence of GIV, or the loss of either of its two functions impaired DNA repair, and induced cell death when challenged with numerous cytotoxic agents. That GIV selectively binds few other BRCT-containing proteins suggests convergent signaling such that heterotrimeric G-proteins may finetune sensing, repair, and outcome after DNA damage.

**GRAPHIC ABSTRACT:** 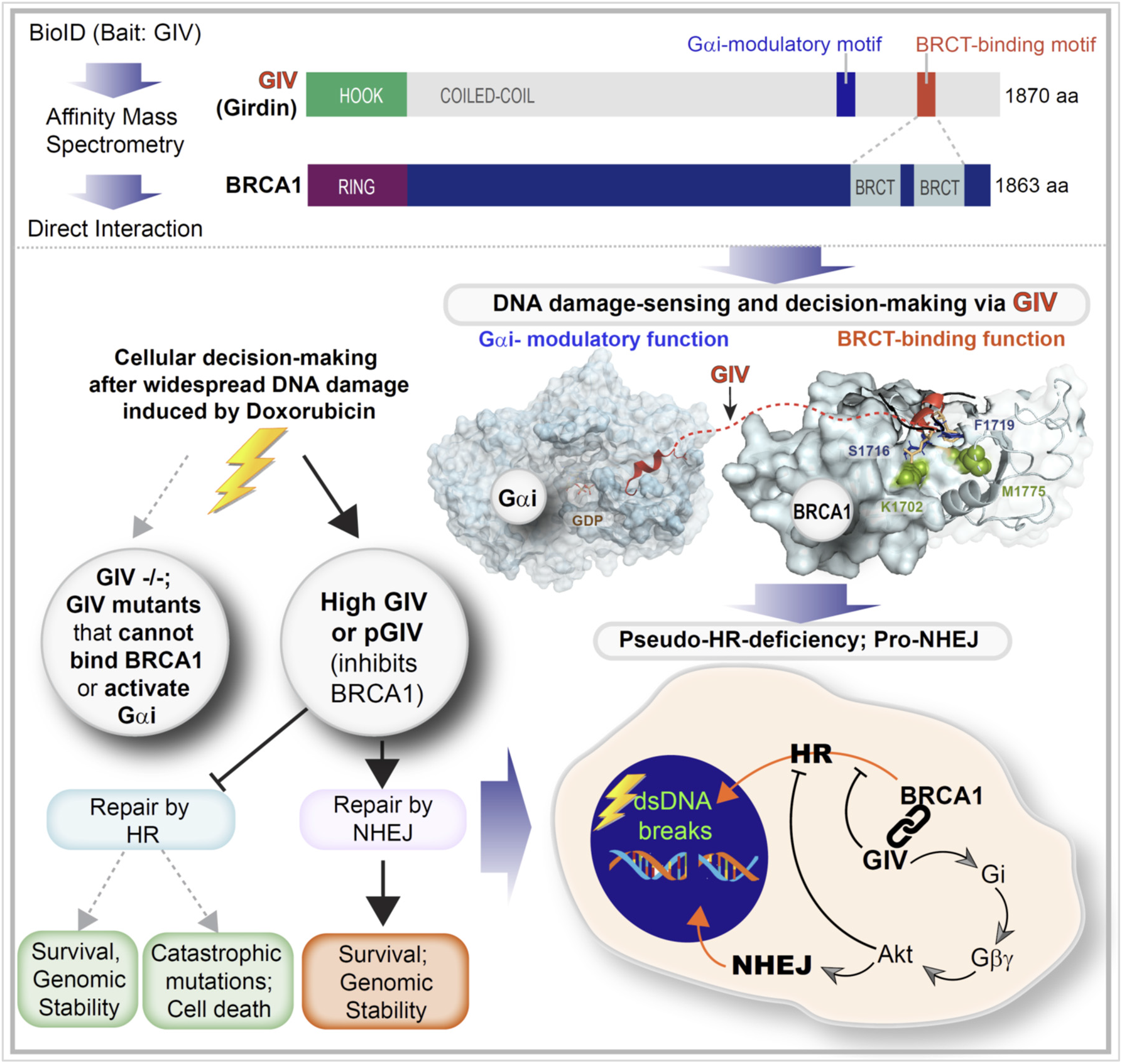

**HIGHLIGHTS:**

- Non-receptor G protein modulator, GIV/Girdin binds BRCA1
- Binding occurs in both canonical and non-canonical modes
- GIV sequesters BRCA1 away from dsDNA breaks, suppresses HR
- Activation of Gi by GIV enhances Akt signals, favors NHEJ

**IN BRIEF:** In this work, the authors show that heterotrimeric G protein signaling that is triggered by non-receptor GEF, GIV/Girdin, in response to double-stranded DNA breaks is critical for decisive signaling events which favor non-homologous end-joining (NHEJ) and inhibit homologous recombination (HR).

## Introduction

Genomic integrity is under constant attack from extrinsic and intrinsic factors that induce DNA damage (*1*). Damaged DNA must be repaired to maintain genomic integrity *via* processes that are personalized and evolved by the cell type, collectively termed as the DNA damage response (DDR) (*2*). DDRs are orchestrated by an incredibly complex network of proteins that sense and assess the type and extent of damage, decide between cell fates (death *vs*. repair), choose a repair pathway, and then initiate and complete the repair process (*3*). For example, the DDR involves the activation of ATM kinase, a member of the phosphoinositide 3-kinase (PI3K)-related protein kinase family (*4*) which is rapidly recruited by the MRE11-RAD50-NBS1 (MRN) complex to chromatin (*5*). Phosphorylation of a large number of substrates follows, which in turn activates cell cycle checkpoints and triggers the recruitment of repair factors to the DSBs. Positive feedback loops are orchestrated to amplify the signals, e.g., ATM phosphorylates the histone variant H2AX (resulting in the formation of the phosphorylated form called γH2AX) (*6*), which recruits additional ATM molecules and further accumulation of γH2AX (*7-9*).

Among the types of DNA damage, DNA double-strand breaks (DSBs) are the most cytotoxic lesions that threaten genomic integrity (*10*). Failure to repair DSBs results in genomic instability and cell death. DNA repair can be achieved by different mean that are commonly grouped into two broad, competing categories (*11*): homologous recombination (HR) and non-homologous end joining (NHEJ) (*12*). HR, which requires a homologous template to direct DNA repair, is generally believed to be a high-fidelity pathway (*13*). By contrast, NHEJ directly seals broken ends; while some believe that repair by NHEJ is imprecise, we have shown that precision can indeed be achieved (*14*). In fact, NHEJ offers an ideal balance of flexibility and accuracy when the damage to DNA is widespread with DSBs featuring diverse end structures (*15*). Consequently, it is believed to represent the simplest and fastest mechanism to heal DSBs (*16*), thus it is the most predominant DSB repair pathway within the majority of mammalian cells. Key molecular players in both pathways have been identified: 53BP1, first identified as a DNA damage checkpoint protein, and Breast cancer type 1 susceptibility protein (BRCA1), a well-known breast cancer tumor suppressor (*17*), are at the center of molecular networks that coordinate NHEJ and HR, respectively.

How the choice of DSB repair pathway is determined at a molecular level has been the subject of intense study for a decade (*18, 19*). Here we reveal a previously unforeseen determinant of the choice of DNA damage, GIV (Girdin), which is a non-receptor activator of heterotrimeric (henceforth, trimeric) G-protein, Gi (*20, 21*). Trimeric G-proteins are a major signaling hub in eukaryotes that gate signaling downstream of 7-transmembrane (7TM)-receptors called GPCRs, and the GPCR/G-protein pathway is of paramount importance in modern medicine, serving as a target of about 34% of marketed drugs (*22*). Although peripheral players in the GPCR/G-protein pathway have been found to have indirect impact on DDR [reviewed in (*23*)], the role of G-proteins in DDR has never been established. Unlike GPCRs that primarily sense the exterior of the cell, GIV-GEM the prototypical member of a family of cytosolic guanine-nucleotide exchange modulators (GEMs), senses and coordinates cellular response to intracellular events (e.g., autophagy, ER-stress, unfolded protein response, inside-out signaling during mechanosensing, etc) by activating endomembrane localized GTPases (*24, 25*). By virtue of its ability to coordinate multiple cellular processes, many of which impart aggressive traits to tumor cells, GIV has emerged as a *bona-fide* oncogene that supports cancer cell stemness, emergence of chemoresistance and invasion, favors aggressive tumor phenotypes in diverse types of cancers, and drives poor survival utcomes [reviewed in (*26*)]. We provide mechanistic insights into how GIV-GEM inhibits HR and concomitantly enhances NHEJ. In doing so, this work not only reveals another pro-tumorigenic role of GIV, but also begins to unravel how endomembrane G-protein signaling shapes decision-making during DDR.

## Materials and Methods

All methods and a Table of reagents, resources, cell lines, equipment and software programs are detailed in *Supplementary Online Materials*. Key methods are briefly mentioned here.

### Biotin Proximity Labeling

BioID was performed as previously described (*27*). Briefly, HEK293T were plated 24 hrs prior to transfection with mycBirA-tagged GIV construct. Biotinylated complexes were then eluted using sample buffer containing excess biotin and heating at 100°C. Prior to mass spectrometry identification, eluted samples were run on SDS-PAGE and proteins were extracted by in gel digest (with trypsin) followed by ultra-high-pressure liquid chromatography (UPLC) coupled with tandem mass spectroscopy (LC-MS/MS) using nano-spray ionization. Identified proteins by mass spec. anaylsis unique to plus biotin samples, but not in minus biotin samples, were analyzed using DAVID and functional annotation was grouped by molecular function and cellular component for GO analysis. Classification with p-value less than 0.05 were considered as significant.

### GIV CRISPR/Cas9 Gene Editing and Validation

Pooled guide RNA plasmids (commercially obtained from Santa Cruz Biotechnology; Cat# sc-402236-KO-2) were used. These CRISPR/Cas9 KO plasmids consists of GFP and girdin-specific 20 nt guide RNA sequences derived from the GeCKO (v2) library and target human Girdin exons 6 and 7. DLD1 parental and GIV KO lines were generated and validated as described before (*28*).

### Long Amplicon PCR

Genomic DNA extraction was performed using the genomic-tip 20/G kit (Qiagen, Cat no. 10223, with corresponding buffer sets) per the manufacturer’s directions. Gene-specific LA-qPCR analyses for measuring DNA damage were performed using Long Amp Taq DNA polymerase (New England Biolabs, Cat no MO323S). Since amplification of a small region would be independent of DNA damage, a small DNA fragment for each gene was also amplified to normalize the amplification of large fragments. Details of PCR conditions are mentioned in the Supplementary Online Materials. The amplified products were then visualized on gels and quantitated with an ImageJ automated digitizing system (National Institutes of Health) based on three independent replicate PCRs. The extent of damage was calculated.

### Stable cell lines with p53BP1 fluorescent reporter

We obtained a lentiviral vector expressing a truncated version of p53BP1 fused to mApple from Dr. Ralph Weissleder, Massachusetts General Hospital, USA (Addgene #69531) (*29*). To establish MDA-MB-231 and HeLa parental and GIV knockout cells stably expressing the reporter, we produced recombinant lentiviruses and transduced each cell line and selected them using Puromycin as described previously (*30*).

### Image acquisition and analysis

We seeded cells in 6- or 96-well glass bottom plates (P-96-1.5H-N or P06-1.5H-N, Cellvis, Mountain View, CA, USA) at densities of 1.25 × 105 or 5 × 103 cells/well, respectively, in imaging base medium (FluoroBrite DMEM media (A1896701, ThermoFisher Scientific, Waltham, MA USA), 1% GlutaMax, 1% PenStrep, 1% sodium pyruvate, and 10% FBS (HyClone)). We performed imaging studies, before and after challenge with various cytotoxic drugs, with an EVOS M7000 Imaging System (ThermoFisher), 40X objective, and the RFP cube for the instrument.

### Homology modeling

The prediction the protein and GIV peptide docked interface was performed by using CABSDOCK (*31*) web server and the final representation was using Pymol visualization tool (*32*). For the analysis the PDB structures 1T19, 1t2V, 1N5O were taken into consideration. 10 residues of GIV “SLSVSSDFLGKD” for their secondary structure prediction was tested on PSIPRED (*33*). The visualization of PDB:1T29 and PDB:1N5O structures was done using MolSoft LLC (https://www.molsoft.com/).

### Statistical Analysis and Replicates

All experiments were repeated at least three times, and results were presented either as average ± SEM. Statistical significance was assessed using one-way analysis of variance (ANOVA) including a Tukey’s test for multiple comparisons. *p < 0.05, **p < 0.01, ***p < 0.001, ****p < 0.0001.

## Results and Discussion

### Proteomic studies suggest a putative role for GIV during DNA damage response

The interactome of a protein dictates its localization and cellular functions. To map the landscape of GIV’s interactome we carried out proximity-dependent biotin identification (BioID) coupled with mass spectrometry (MS) (**Fig 1A**). BirA-tagged GIV construct was validated by immunoblotting and found to be expressed in cells as full-length proteins of expected molecular size (∼250 kDa) (**Fig 1B**). Samples were subsequently processed for protein identification by Mass Spectrometry. Gene ontology (GO) cellular component analysis, as determined by DAVID GO, revealed that GIV-proximal interactors were in both cytoplasmic and nuclear compartments (**Fig 1C;** See **Supplementary Extended Data 1**); one interactor (i.e., BRCA1) was predicted to bind GIV across three different cellular compartments (**Fig 1C**; red bars). GO-molecular function analysis revealed that “DNA binding proteins” was the most enriched class of proteins in GIV’s interactome (**Fig 1D; Supplementary Extended Data 2**). BRCA1 was a notable interactor within that class, and the serine/threonine-specific protein kinase, Ataxia telangiectasia and Rad3 related protein (ATR), that coordinates DNA damage sensing and repair was another (**Fig 1E**). A reactome pathway analysis of DNA-binding proteins in the GIV’s interactome showed that GIV’s interactome enrichment for proteins that participate in gene transcription, regulation of cell cycle, and in DNA repair (**Fig 1E-F**). These findings were rather surprising because GIV’s presence on nuclear speckles was described almost a decade ago (*34*), and yet, among all the functions of GIV that have emerged since its discovery in 2005, little to nothing is known about GIV’s role in sensing/signaling during intranuclear processes.

**Figure 1.**
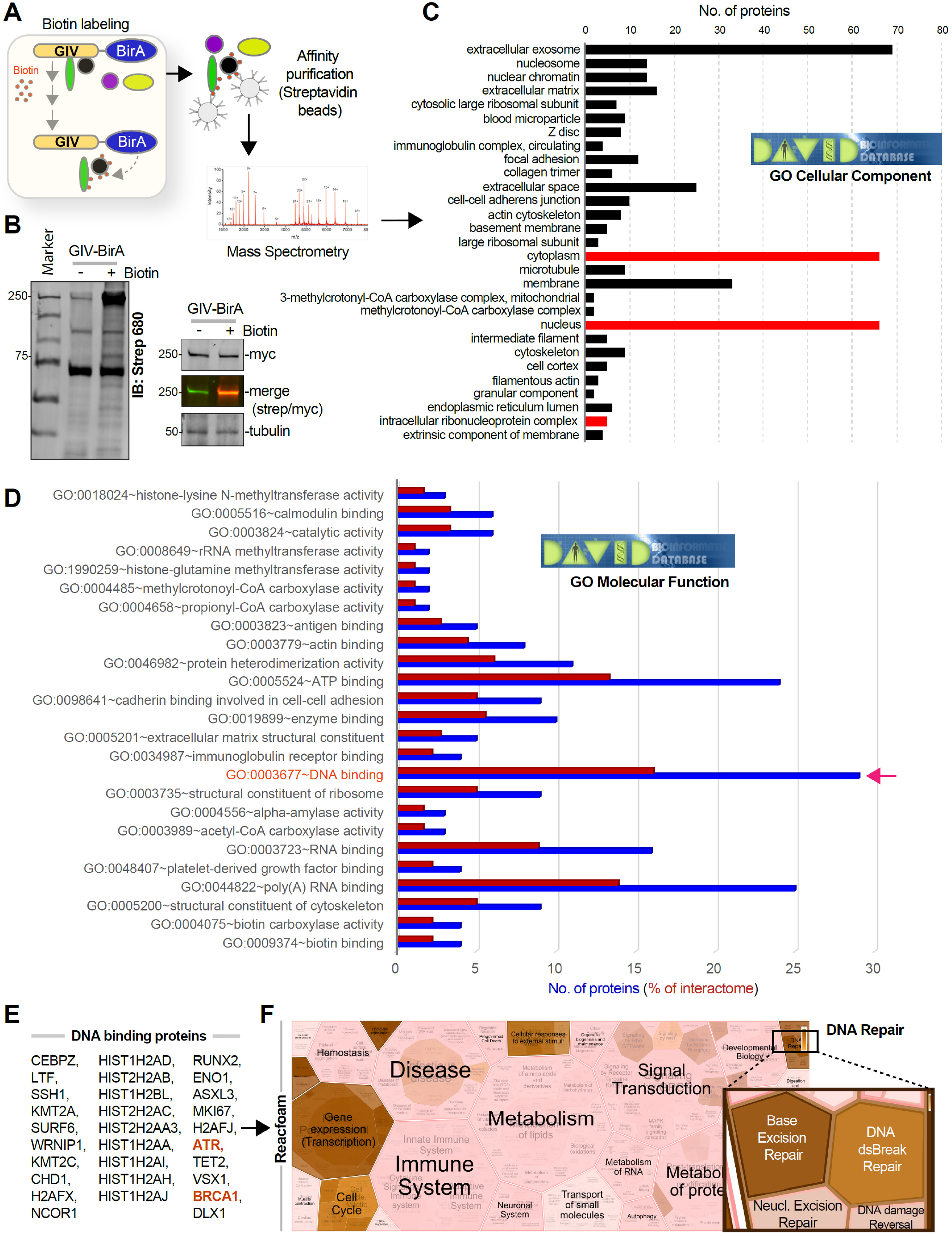
Proteomic studies suggest an intranuclear role of GIV/Girdin in DNA damage repair response. **A**. Schematic outlining key steps in BioID studies to identify the GIV interactome. **B**. Immunoblots confirm biotinylation in HEK whole cell lysates (left) and expression of the BirA-tagged full-length GIV construct as a protein of expected size (right). **C-D**. Bar plots show GO analyses [cellular component (C) and molecular function (D)] for bioID-identified GIV interactome. Red bars in C indicate putative compartments where GIV binds BRCA1. Blue and red bars in D indicate total number of interacting proteins and % representation, respectively. Red arrow in D indicates the molecular function category where BRCA1 was identified. **E-F**. DNA-binding proteins (listed in E) that were identified in GIV’s interactome were analyzed by Reactome.org and visualized as hierarchical reacfoam (in F). Inset in top right corner is magnified to highlight the overrepresentation of DNA repair pathways.

### GIV is required for DNA damage response

We generated HeLa cells without GIV using CRISPR Cas9 and subsequently exposed them to Doxorubicin followed by several commonly used readouts of DDR (**Fig 2A**; **S1A-B**). A mixture of -/- (henceforth, GIV KO) clones were pooled to recapitulate the clonal heterogeneity of parental HeLa cells (**Fig S1B**), and near-complete depletion of GIV (estimated ∼95% by band densitometry) was confirmed by immunoblotting (**Fig 2B**). We chose HeLa cells because DDR has been extensively studied in this cell line (*35, 36*) and because HeLa cells have defective p53 (*37*). The latter is relevant because GIV/CCDC88A aberrations (gene amplification) co-occur with defects in the tumor suppressor TP53 (TCGA pancancer profile; cbioportal.org); ∼36% of tumors with aberrant CCDC88A expression was also associated with mis-sense and truncating driver mutations in TP53. We chose Doxorubicin (henceforth, Dox) for inducing DNA damage because it is a widely used anthracycline anticancer agent and its impact on DNA integrity in HeLa cells has been mapped for each cell cycle with demonstrated reproducibility (*38*). Compared to parental cells, fewer metabolically active GIV KO cells survived after a Dox challenge, as determined using a MTT assay (**Fig 2C; Fig S1C**), indicating that in the absence of GIV, cells show markedly reduced survival from cytotoxic lesions induced by Dox. GIV KO cells showed increased susceptibility also to two other cytotoxic drugs, Cisplatin and Etoposide (**Fig S1D-E**). The lower IC50 values in the case of GIV KO cell lines for all 3 drugs (**Fig 2C**) imply that GIV is required for surviving cytotoxic lesions induced by the most commonly used cytotoxic drugs.

**Figure 2.**
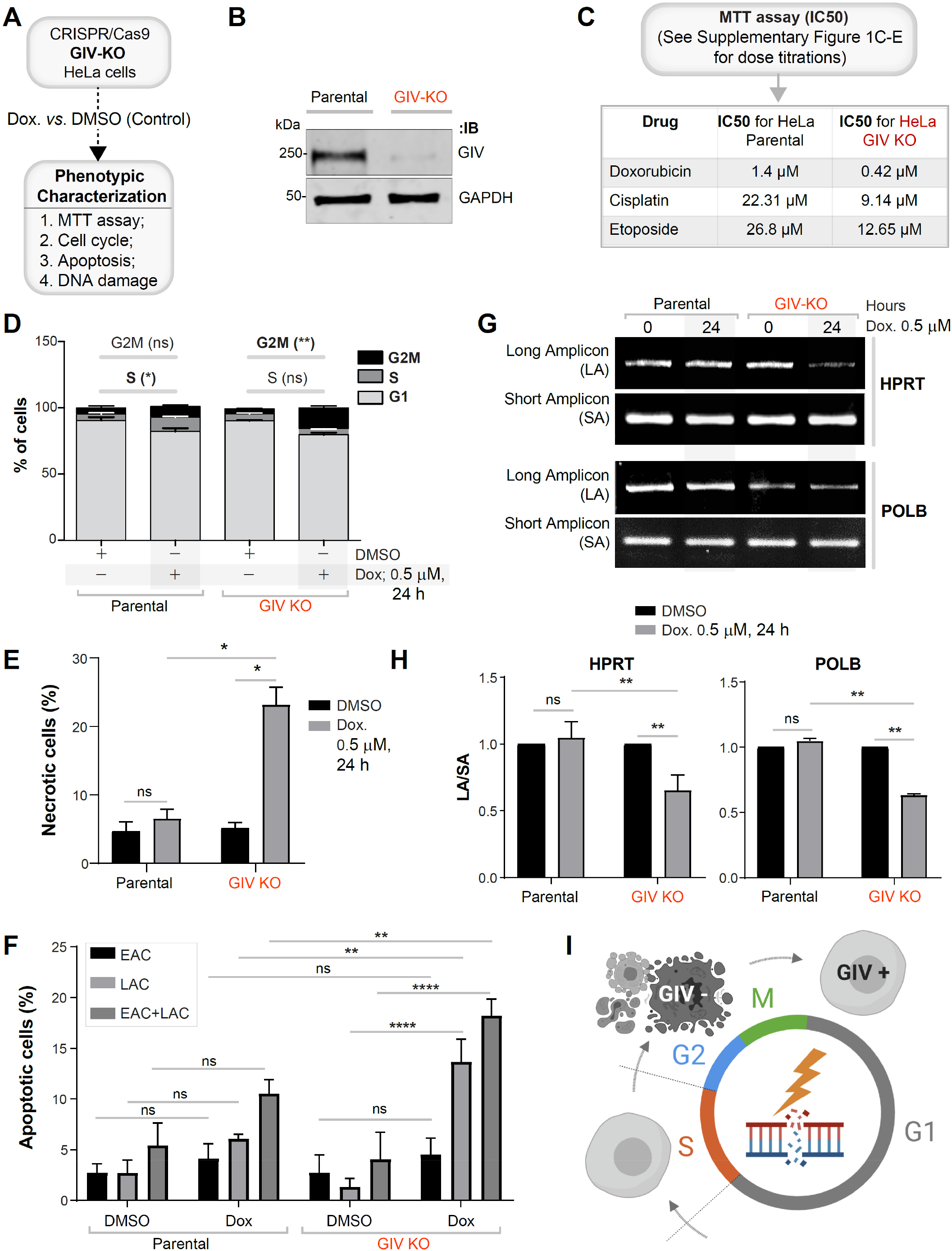
DNA damage repair response is impaired in cells without GIV. **A**. Schematic outlining the cell lines and phenotypic assays displayed in this figure. **B**. Immunoblot of GIV-depleted (by CRISPR Cas9) and control (Parental) HeLa cell lysates showing depletion of full-length endogenous GIV. See also Fig S1 for how pooled KO lines were generated. **C**. Table of IC50 values for 3 different drugs tested on parental and GIV KO HeLa cells, as determined using MTT assays. See **Fig S1C-E** for the dose-dependent survival curves. **D**. Stacked bar graphs showing the percentage of cells at various stages of the cell cycle (G1, S and G2/M) after challenged with Dox or vehicle control (DMSO). Histograms are shown in **Fig S1F**. Data displayed as mean ± S.E.M. and one-way ANOVA using Tukey’s multiple comparisons test was used to determine significance. (*; p ≤ 0.05, **; p ≤ 0.01; ns = not significant). **E-F**. Bar graphs display the % necrotic (E) or apoptotic (early, EAC; late, LAC; or combined) cells after challenged with either Dox or vehicle control (DMSO), as assessed by annexin V staining and flow cytometry. See **Fig S1G** for the dot plot diagrams. **G-H**. Long amplicon qPCR (LA-QPCR) was used to evaluate genomic DNA SB levels in control vs. GIV KO cells. Representative gel showing PCR-amplified fragments of the *HPRT* (G, top panel) and *POLB* (G, bottom panel) genes. Amplification of each large fragment (upper panels) was normalized to that of a small fragment of the corresponding gene (bottom panels) and the data were expressed as lesion frequency/10 Kb DNA and displayed as bar graph in H. Full-length gels can be seen in **Fig S1H**. Data displayed as mean ± S.E.M. and one-way ANOVA to determine significance. (**; p ≤ 0.01; ****; p ≤ 0.0001; ns = not significant). **I**. Schematic summarizing the findings in cells with (parental; GIV +) or without GIV (GIV KO; GIV -).

Because cell cycle is a key determinant of the choice of repair pathway, next, we asked if GIV may impact one or more of the three checkpoints (G1/S, S phase, and G2/M) where cell cycle may be arrested in response to DNA damage. We found that Dox-challenged parental cells, as expected for cells with defective p53, escaped the G1/S checkpoint (*39*), and instead, preferentially showed arrest in S/G2 phase; however, GIV KO cells showed no such S phase arrest and instead arrested in the G2 phase (**Fig 2D**; **Fig S1F**). Because chromosome duplication occurs during the “S phase” (the phase of DNA <u>s</u>ynthesis) and this phase surveys DNA for replication errors (*40*), failure of GIV KO cells to arrest in S phase indicates that this “checkpoint” is impaired (i.e., bypassed). Because irreparable DNA injury leads to the accumulation of mutations, which in turn may induce either apoptosis and necrosis (*41*), next we analyzed cell death by flow cytometry using a combination of annexin V and propidium iodide (PI) staining. Compared to parental control cells, Dox challenge induced a significantly higher rate of cell death in GIV KO cells (**Fig 2E-F**; **Fig S1G**), *via* both necrosis (**Fig 2E**) and apoptosis (specifically, late apoptosis; **Fig 2F**).

To examine whether higher cell death was related to impaired repair activity and an accumulation of DNA strand breaks in GIV KO cells, genomic DNA was isolated from parental and GIV KO cells, with or without Dox challenge and the levels of strand breaks in the *HPRT* and *POLB* genes were compared using long amplicon quantitative PCR (LA-qPCR) as described previously (*42*). Strand breaks were measured for both the genes using a Poisson distribution, and the results were expressed as the lesion/10 kb genome (*43*). A decreased level of the long amplicon PCR product (12.2 kb of the POLB or 10.4 kb region of the HPRT gene) would reflect a higher level of breaks; however, amplification of a smaller fragment for each gene is expected to be similar for the samples, because of a lower probability of breaks within a shorter fragment. A higher level of DNA lesion frequency was observed per 10 Kb in the genomic DNA of GIV KO cells than in the DNA of parental controls (**Fig 2G-H**; **Fig S1H**), indicating a role of GIV in DNA repair.

Reduced cell survival (**Fig 2C**), cell cycle arrest (**Fig 2D**), higher cell death (**Fig 2E-F**) and the accumulation of cytotoxic lesions (**Fig 2G-H**) in GIV KO cells was also associated with reduced growth in anchorage-dependent clonogenic growth assays (**Fig S1I**).

To test if the pro-survival functions of GIV in the setting of cytotoxic lesions is cell-type specific, we compared 2 other cell lines, the MDA-MB-231breast and DLD-1 colorectal cancer lines (**Fig S2A**). We generated GIV KO MDA-MB-231 cell lines using CRISPR Cas9 (see validation in **Fig S2B-C**) and used the previously validated GIV KO DLD-1 cells (*28*). We exposed these cells to Doxorubicin. Survival was significantly impaired in all the GIV KO cell lines (**Fig S2D-F**), implying that our findings in HeLa cells may be broadly relevant in diverse cancers.

Taken together, these findings demonstrate that GIV is required for DNA repair; in cells without GIV, cell survival is reduced, S phase checkpoint is lost, and DNA repair is impaired, leading to the accumulation of mutations in *POLB* and *HPRT* (see **Fig 2I**). These findings suggest that genotoxic insult in the absence of GIV may lead to the accumulation of catastrophic amounts of mutations that may ultimately trigger cell death.

### The C-terminus of GIV binds tandem BRCT modules of BRCA1

We next sought to validate the major BioID-predicted interaction of GIV, i.e., BRCA1. To determine if GIV and BRCA1 interact in cells, we carried out coimmunoprecipitation (Co-IP) assays and found that the two full-length endogenous proteins exist in the same immune complexes (**Fig 3A**). BRCA1 features two prominent modules that mediate protein-protein interactions, an N-terminal RING domain, which functions as an E3 ubiquitin ligase (*44*), and a C-terminal BRCT repeat domain, which functions as phospho-protein binding module (*45*). Pulldown assays using recombinant GST-tagged BRCA1-NT (RING) or CT (tandem BRCT repeats) proteins immobilized on Glutathione beads and lysates of HEK cells as a source of FLAG-tagged GIV showed that full-length GIV binds BRCT, but not the RING module (**Fig 3B**). We noted that GIV is predicted to also interact with other BRCT-domain containing DDR pathway proteins, e.g., DNA Ligase IV (LIG4) and Mediator Of DNA Damage Checkpoint 1 (MDC1) [Human cell map, cell-map.org; a database of BioID proximity map of the HEK293 proteome; accessed on 01/06/2020] and with BARD1 [BioGRID, thebiogrid.org; accessed 09/05/2020]. Pulldown assays with these BRCT modules showed that GIV bound DNA Ligase and BARD1, but not MDC1 (**Fig 3B**), suggesting that while GIV can promiscuously bind multiple DDR pathway proteins that contain the BRCT module, there may be a basis for selectivity within such apparent promiscuity. As a positive control for BRCT-binding protein, we tracked by immunoblotting the binding of BACH1 from the same lysates, which bound BRCA1’s tandem BRCT module, as expected (*46*), and to a lesser extent with DNA Ligase (**Fig 3B**).

**Figure 3.**
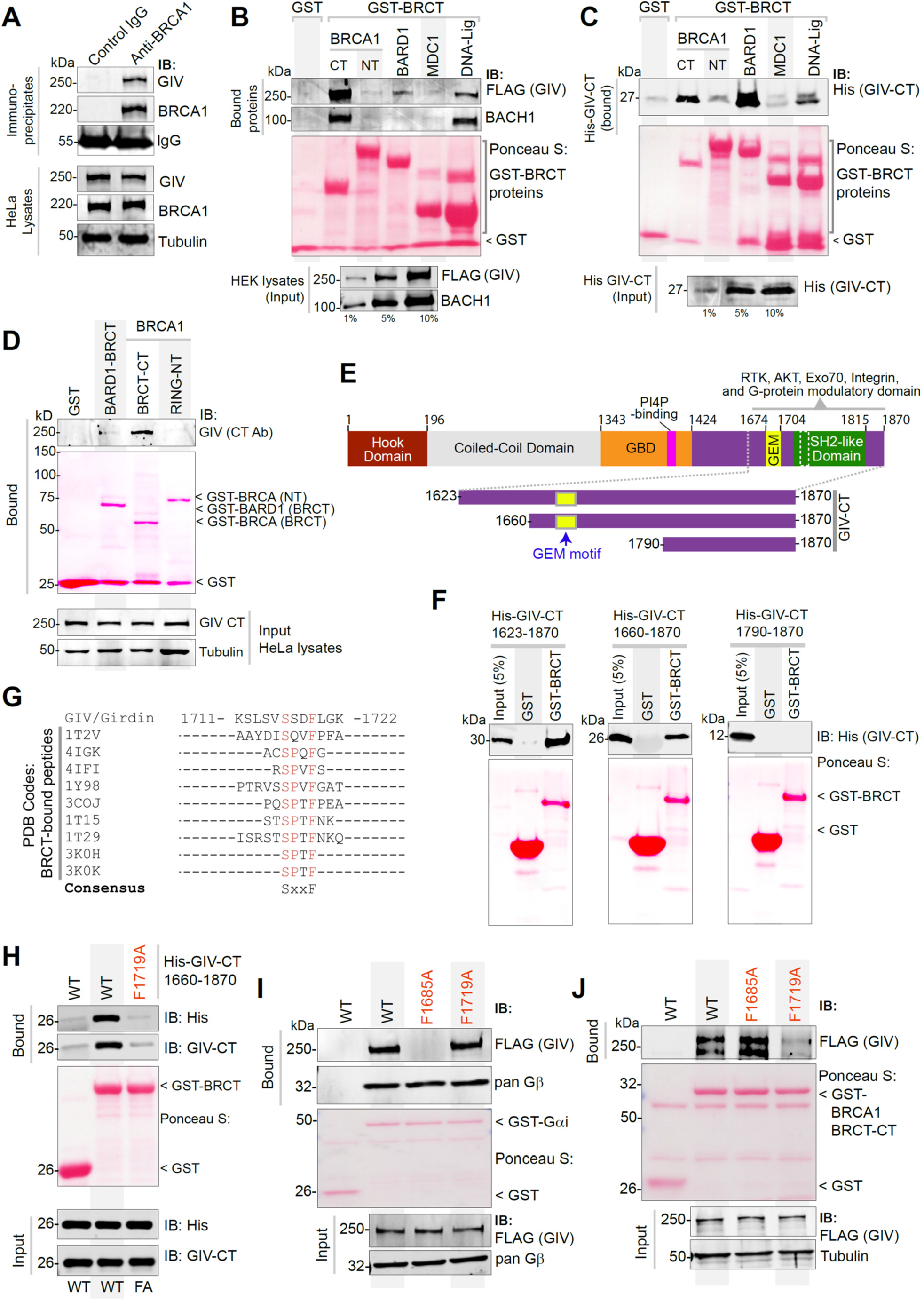
GIV directly binds BRCT module of BRCA1. **A**. Coimmunoprecipitation assays were carried out on lysates of HeLa cells using anti-BRCA1 antibody or control IgG and immune complexes (top) and lysates (bottom) were analyzed for GIV and BRCA1 by immunoblotting. **B**. Lysates of HEK cells exogenously expressing FAG-tagged full-length GIV were used as the source of GIV and BACH1 (positive control for known BRCA1-binding protein) in pulldown assays with GST-tagged BRCA1 fragments and BRCT modules of various indicated proteins (visualized using Ponceau S). Bound proteins (top) and lysates (bottom) were analyzed for GIV and BACH1. **C**. Pulldown assays were carried out using recombinant His-GIV-CT (aa 1660-1870) and GST-BRCT modules as in B. Bound GIV was visualized by immunoblotting (anti-His). **D**. Pulldown assays were carried out using lysates of HeLa cells as the source of endogenous full-length GIV with GST-BRCA1 and BARD1. Bound GIV was visualized by immunoblotting. See also **Fig S3** for similar studies with Cos7 and Hs578T cell lysates. **E-F**. Recombinant GIV-CT proteins of various lengths (see schematic E) were used in pulldown assays with GST-BRCT module of BRCA1. Bound GIV-CT fragments were analyzed in F by immunoblotting (His). **G**. Alignment of GIV’s C-terminal sequence with known phosphopeptides that bind BRCA1, as confirmed by x-ray crystallography (PDB codes on the left). The consensus SxxF sequence is shown (evolutionary conservation of the SxxF motif and its relationship with other motifs on GIV-CT is shown in **Fig S4**). **H**. Pulldown assays were carried out using His-GIV-CT WT or F1719A mutant with GST/GST-BRCA1 and bound GIV was analyzed by immunoblotting. **I-J**. Pulldown assays were carried out with either GDP-loaded GST-Gαi3 (I) or GST-BRCA1 (BRCT; J) proteins and lysates of HEK cells exogenously expressing FLAG-tagged GIV wild-type (WT) or GIV mutants that do not bind G protein (F1685A) (*20*) or do not bind BRCA1 (F1719A; current work). Bound proteins were visualized by immunoblotting using anti-FLAG IgG.

We next asked if the C-terminus of GIV can directly bind BRCA1; we focused on GIV’s C-terminus (GIV-CT) because numerous studies have underscored the importance of GIV-CT as an unstructured and/or intrinsically disordered domain that scaffolds key proteins within major signaling cascades to mediate dynamic pathway crosstalk (*47, 48*). GST pulldown assays using recombinant His-tagged GIV-CT^1660-1870^ and various GST-DDR pathway proteins showed that GIV’s CT is sufficient to bind the C-terminal tandem BRCT domain of BRCA1 (**Fig 3C**). Because we used purified recombinant proteins in this assay, we conclude the GIV•BRCA1 interaction observed in cells is direct. Using lysates from multiple different cell types as source of GIV (Hs578T, **Fig 3D**; Cos7 and HeLa; **Fig S3A-B**) we further confirmed that endogenous full-length GIV binds the C-terminal tandem BRCT domain of BRCA1 (but not its RING domain) and weakly with BARD1.

Domain-mapping efforts, using various fragments of GIV-CT (aa 1623-1870, 1660-1870, and 1790-1870; **Fig. 3E**) helped narrow the region within GIV that binds BRCA1. The longer GIV-CT fragments bound, but the shortest fragment (1790-1870) did not (**Fig. 3E-F**), indicating that the sequence of GIV that lies between aa 1660-1790 could be the key determinant of binding. A sequence alignment of this region on GIV against known interactors of BRCA1’s tandem BRCT repeats revealed the presence of a canonical BRCT-binding phospho-peptide sequence of the consensus “phosphoserine (pSer/pS)-x-x-Phenylalanine (Phe; F)” (**Fig 3G**). The structural basis for such binding has been resolved (*49*). This newly identified putative BRCT-binding motif in GIV had three notable features: First, this motif (^1716^SSDF^1719^) is distinct from and farther downstream of GIV’s Gαi-modulatory motif (31 aa ∼1670-1690) (**Fig S4A**), suggesting that they may be functionally independent. Second, the SxxF motif is evolutionarily conserved in higher vertebrates (birds and mammals) (**Fig S4A**), suggesting that GIV could be a part of the complex regulatory capacities that evolved later (*50*). Third, multiple independent studies have reported that the Ser in ^1716^SSDF^1719^ is phosphorylated (**Fig S4B**), suggesting that GIV•BRCA1 complexes may be subject to phosphomodulation. Site-directed mutagenesis that destroys the consensus motif (by replacing Phe with Ala; F1719A) resulted in a loss of binding between GIV-CT and BRCA1 (**Fig 3H**), thereby confirming that the putative BRCT-binding motif is functional and implicating it in the GIV•BRCA1 interaction. The independent nature of the BRCA1-binding and Gαi-modulatory motif was confirmed in pulldown assays with full-length WT and mutant GIV proteins (**Fig 3I-J**); the BRCA1 binding-deficient F1719A mutant protein selectively lost binding to GST-BRCA1, but not GST-Gαi3, and the well-characterized G-protein binding-deficient F1685A mutant protein (*20, 21*) selectively lost binding to GST-Gαi3, but not GST-BRCA1.

Collectively, these findings demonstrate that GIV binds BRCA1 via its C-terminally located BRCT-binding motif. This motif is sensitive to disruption *via* a single point mutation but specific enough that such mutation does not alter GIV’s ability to bind Gαi-proteins.

### GIV binds BRCA1 in both phospho-dependent and -independent modes via the same motif

We next asked how GIV binds BRCA1(BRCT). BRCT modules are known to bind ligands *via* two modes—(i) canonical, phospho-dependent (e.g., BACH1, CtIP, Abraxas) and (ii) non-canonical, phospho-independent (e.g., p53) (*51*); while the structural basis for the former has been resolved (*49*), the latter remains unclear. Because bacterially expressed His-GIV-CT directly binds the tandem BRCA1-BRCT (**Fig 3C**), the GIV•BRCA1 (BRCT) interaction appears phospho-independent. As positive controls for canonical phospho-dependent binding, we used BACH1 and CtIP, two *bona fide* binding partners of BRCA1-BRCT module. Recombinant His-GIV-CT did not impact the canonical mode of binding of either BACH1 (**Fig 4A**) or CtIP (**Fig 4B**) to BRCA1-BRCT, suggesting that unphosphorylated GIV binds BRCA1 at a site that is distinct from the interdomain cleft where BACH1 or CtIP are known to occupy (*49*). Furthermore, binding of GIV to the tandem BRCT was enhanced ∼3-5-fold in the presence of the most frequently occurring mutation in BRCA1, M1775R (**Fig 4C**); this mutation is known to abrogate canonical mode of phosphopeptide binding by destroying a hydrophobic pocket that otherwise accommodates the Phe in the pSxxF consensus (see **Fig S5A**) (*52*). The unexpected increase in binding to the BRCA1-M1775R mutant was also observed in the case of p53, which is another phospho-independent BRCA1(BRCT)-interacting partner (*53*) (**Fig 4D**). The expected disruptive effect of this mutation could, however, be confirmed in the case of both BACH1 (**Fig S5B**) and CtIP (**Fig S5C**). These findings demonstrate that GIV binds BRCA1 *via* a non-canonical phospho-independent mechanism that is distinct from CtIP and BACH1.

**Figure 4.**
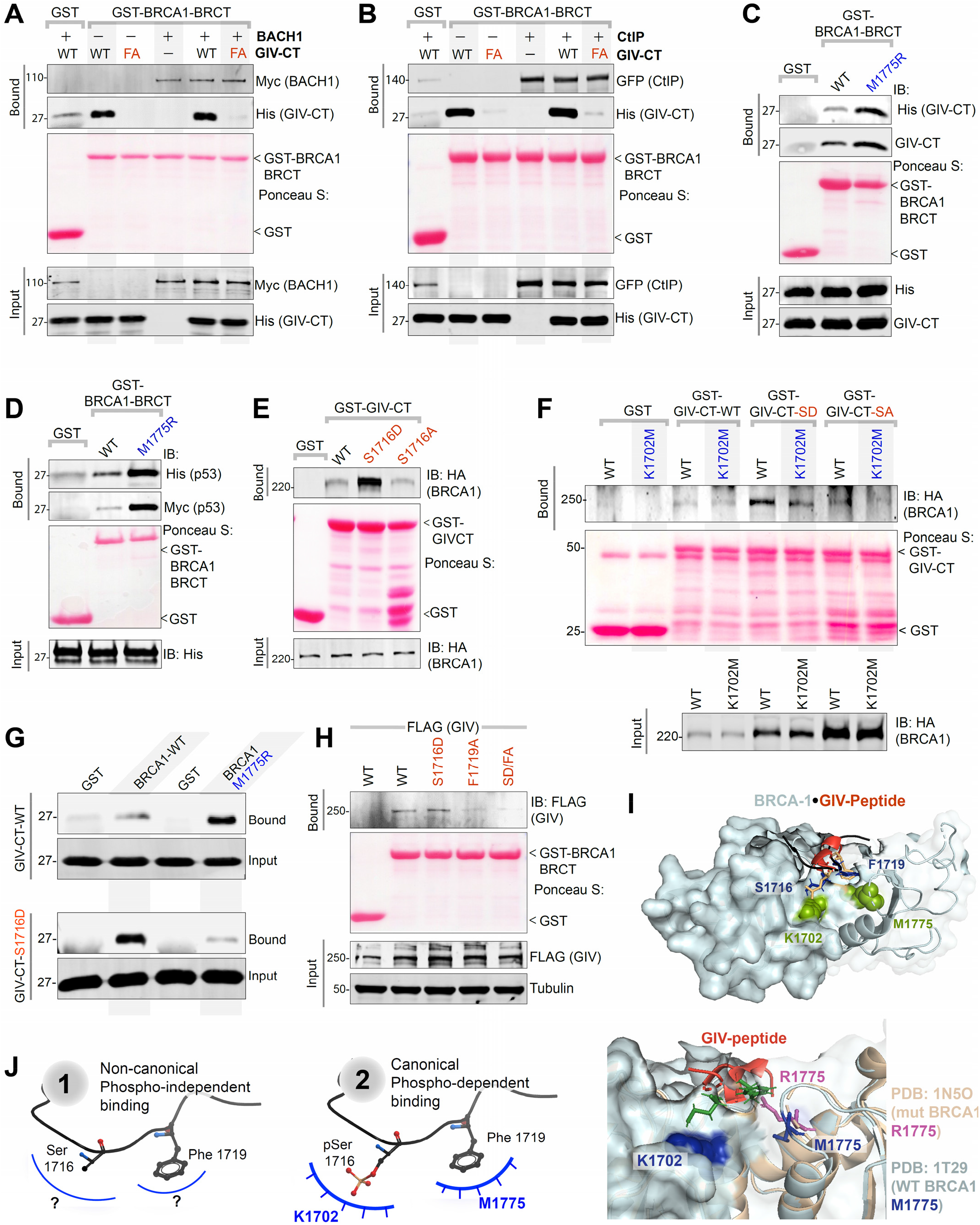
GIV binds BRCA1 *via* both canonical (phosphodependent) and non-canonical (phosphoindependent) mechanisms. **A-B**. Binding of unphosphorylated GIV with BRCA1 does not compete with canonical, phospho-dependent binding of BACH1 (A) or CtIP (B). Pulldown assays were carried out using lysates of HEK cells as source of myc-BACH1 (A) or GFP-CtIP (B) and recombinant GST/GST-BRCA1 proteins, in the presence (+) or absence (-) of either wild-type (WT) or BRCA-binding deficient F1719A (FA) mutant His-GIV-CT. Bound proteins were visualized by immunoblotting with anti-His (GIV), anti-myc (BACH1; A), or anti-GFP (CtIP; B) IgGs. See also **Fig S5**. **C-D**. Pulldown assays were carried out using His-GIV-CT (C) or His-TP53 (D) and GST or GST-BRCA1 (WT and M1775R mutants). Bound proteins were visualized by immunoblotting with anti-His IgG. **E**. Pulldown assays were carried out using lysates of HEK cells as the source of HA-BRCA1 (full length) with either GST (control) or wild-type (WT) and phosphomimic (S1716D) or non-phosphorylatable (S1716A) mutant GST-GIV-CT. Bound BRCA1 was visualized by immunoblotting. **F**. Pulldown assays were carried out as in E, using lysates of HEK cells exogenously expressing either wild-type (WT) or K1702M mutant of HA-BRCA1. **G**. Pulldown assays were carried out using recombinant His-GIV-CT (WT or S1716D) and either GST-BRCA1 WT or M1775R mutant protein as in C. Bound GIV was visualized by immunoblotting using anti-His IgG. **H**. Lysates of HEK cells exogenously expressing full-length GIV-FLAG constructs were used as the source of GIV in pulldown assays with GST/GST-BRCA1. Bound GIV was visualized using anti-FLAG IgG. **I**. Homology model of phospho-dependent GIV•BRCA1 complex (I; *top*) built using the solved crystal structure of BACH1•BRCA1 complex (PDB: IT29) as a template. GIV = red; major residues on BRCA1 or GIV that were mutated here are labeled. Impact of M1775R mutant BRCA1 posing a steric clash with F1719 (GIV) is highlighted (I; *bottom*). **J**. Schematic summarizing the two modes of binding of the same ^1716^SxxF^1719^ sequence on GIV-CT to the BRCT module of BRCA1. The structural basis for phospho-independent binding remains unknown (*left*; “?”).

Because ∼10 high-throughput (HTP) studies have confirmed that Ser^1716^ within the BRCA1-binding motif of GIV is phosphorylated (**Fig S5C**), presumably by one of the many DDR and cell-cycle regulatory kinases (**Fig S5D**), we asked if the GIV•BRCA1 interaction is phosphomodulated. Phosphomimic (Ser^1716^→Asp; S1716D) and non-phosphorylatable (Ser^1716^→Ala; S1716A) mutants of GST-GIV-CT were generated, rationalized based on systematic peptide screening studies demonstrating that Glu/Asp-x-x-Phe peptides bind BRCT modules with ∼10-fold higher affinity (*54*). Binding of BRCA1 was accentuated with GIV-S1716D mutant but restored to levels similar to WT in the case of GIV-S1716A mutant (**Fig 4E**), indicating that the GIV•BRCA1 interaction may be phosphoenhanced and that the -OH group in Ser (which is absent in Ala; A) is not essential for the interaction. The phosphate group in the consensus pSxxF mediates polar interactions with S1655/G1656 in β1 and K1702 in α2 of BRCA1 (*49*), and a K1702M mutant has previously been shown to impair phospho-dependent canonical mode of binding (*50*). We found that the observed phosphoenhanced GIV•BRCA1 in **Fig 4E** is virtually abrogated in the case of BRCA1-K1702M (**Fig 4F**), indicating that upon phosphorylation at S1716, GIV may bind BRCA1 in a phospho-dependent canonical mode. Finally, in pulldown assays with the BRCA1-M1775R mutant, binding was inhibited to the phosphomimic GIV-S1716D mutant, but not to GIV-WT (**Fig 4G**), likely *via* the obliteration of the binding pocket for the F1719, as has been reported in the canonical binding mode (*52*). That the F1719 is also important for phospho-dependent binding was also confirmed; the addition of F1719A mutation to S1716D mutation disrupted binding to BRCA1 (**Fig 4H**), indicating that the same BRCA1-binding motif participates in both modes of binding. Homology models of GIV•BRCA1 co-complexes (**Fig 4I**; *top*), built using the solved structure of canonical BACH1•BRCA1 co-complex (PDB:1T29) as template further confirmed that phospho-dependent canonical mode of binding and disrupted binding when M1775 is mutated to R (**Fig 4I**; *bottom*) is compatible with the observed biochemical studies.

Taken together, these findings support the conclusion that GIV binds BRCA1 in two different modes: a non-canonical phospho-independent mode, the structural basis for which remains unknown (**Fig 4J**; *left*), and a canonical phospho-dependent mode (**Fig 4J**; *right*). Both modes of binding occur *via* the same motif in GIV.

### Both GIV•BRCA1 and GIV•Gαi interactions are required for DNA repair

To dissect the role of the GIV•BRCA1 interaction, we rescued GIV KO HeLa cell lines with either GIV-WT or single-point specific mutants of GIV that either cannot bind BRCA1 (F1719A) or cannot bind/activate Gαi-proteins (F1685A) and used them in the same phenotypic assays as before (**Fig 5A**). First, we confirmed by immunoblotting that the G418-selected clones stably express physiologic amounts of GIV-WT/mutants at levels similar to endogenous (**Fig 5B**). When challenged with Dox, cisplatin, or etoposide, survival, as determined using a MTT assay was significantly reduced in the cells expressing either mutant compared to GIV-WT (**Fig 5C; Fig S6A-C**). The lower IC50 values in the case of GIV mutant cell lines for all 3 drugs (**Fig 5C**) imply that both functions of GIV, i.e., BRCA1-binding and G protein-binding/activating, are required for surviving cytotoxic lesions induced by commonly used cytotoxic drugs. Lower survival was associated with G2/M phase arrest in both mutant lines (**Fig 5D**; **S6A**). The S phase checkpoint, however, was intact in cells expressing GIV-WT and GIV-F1685A mutant, but not in GIV-F1719A mutant (**Fig 5D**), indicating that disruption of the GIV•BRCA1 interaction blocks the S phase checkpoint. Flow cytometry studies showed that cell death, both necrosis (**Fig 5E, S6B**) and apoptosis (late apoptosis; LAC; **Fig 5F, S6B**), was significantly increased in both mutant-expressing lines compared to GIV-WT. The extent of death was higher in GIV-F1719A mutant lines, indicating that disruption of the GIV•BRCA1 interaction is catastrophic. Consistently, the burden of mutations was increased in both mutant lines, but to a higher degree in GIV-F1719A mutant lines (**Fig 5G-H**; **S6C**).

**Figure 5.**
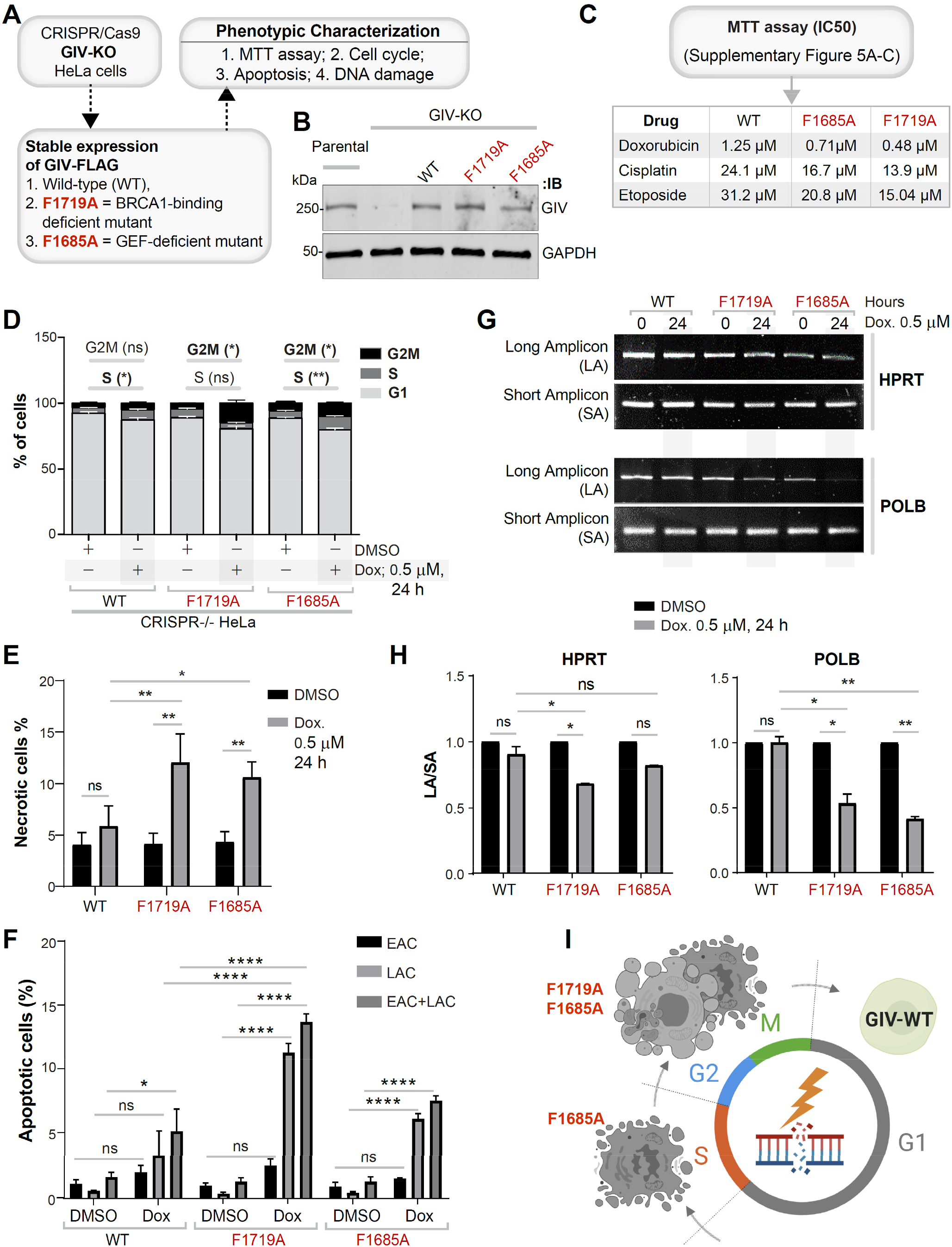
DNA damage repair response is impaired in cells expressing mutant GIV that cannot bind BRCA1 (F1719A) or bind/activate G proteins (F1685A). **A**. Schematic outlining the cell lines and phenotypic assays displayed in this figure. **B**. Immunoblot of the HeLa cell lysates showing depletion of full-length endogenous GIV, followed by rescue WT and mutant GIV at levels close to endogenous. **C**. Table of IC50 values for 3 different drugs tested on parental and GIV KO HeLa cells, as determined using MTT assays. See **Fig S6A-D** for the dose-dependent survival curves. **D**. Stacked bar graphs showing the percentage of cells at various stages of cell cycle (G1, S and G2/M) after challenged with Dox or vehicle control (DMSO). Data displayed as mean ± S.E.M. and one-way ANOVA using Tukey’s multiple comparisons test was used to determine significance. (*; p ≤ 0.05, **; p ≤ 0.01; ns = not significant). Histograms are shown in **Fig S6E**. **E-F**. Bar graphs display the % necrotic (E) or apoptotic (early, EAC; late, LAC; or combined) cells after challenge with either Dox or vehicle control (DMSO) as assessed by annexin V staining and flow cytometry. See **Fig S6F** for the dot plot diagrams. **G-H**. Long amplicon qPCR (LA-QPCR) was used to evaluate genomic DNA SB levels in various HeLa cell lines. Representative gel showing PCR-amplified fragments of the *HPRT* (G, top panel) and *POLB* (G, bottom panel) genes. Amplification of each large fragment (upper panels) was normalized to that of a small fragment of the corresponding gene (bottom panels) and the data were expressed as lesion frequency/10 Kb DNA and displayed as bar graph in H. Full length gels can be seen in **Fig S6G**. Data displayed as mean ± S.E.M. and one-way ANOVA to determine significance. (*; p ≤ 0.05; ****; p ≤ 0.0001; ns = not significant). **I**. Schematic summarizing the findings in cells with GIV-WT or mutants that either cannot bind G protein (F1685A) or BRCA1 (F1719A).

Taken together, these results demonstrate that both functions of GIV (BRCA1-binding and Gαi binding and activation) are important for GIV’s role in mounting a DDR. The use of GIV KO cell lines rescued with WT or specific binding-deficient mutants further pinpointed the role of each function in the process (**Fig 5I;** summarized in **Supplementary Table 1**). The GIV•BRCA1 interaction was required for S phase checkpoint arrest, cell survival, and DNA repair. However, GIV’s Gαi-modulatory function was somehow important for cell survival and the efficiency of DNA repair.

### GIV inhibits HR and favors NHEJ

We next asked how GIV’s ability to bind BRCA1 or activate Gαi might impact the choice of the pathway for DNA repair. While γH2AX is responsible for the recruitment of many DNA maintenance and repair proteins to the damaged sites, including 53BP1 and RAD51 (*55*), the preferential accumulation of 53BP1 indicates NHEJ, whereas the preferential accumulation of Rad51 indicates HR (*12*) (**Fig 6A**). BRCA1 favorably activates Rad51-mediated HR repair and actively inhibits 53BP1-mediated NHEJ repair (*56*). We found that nuclear accumulation of 53BP1, as determined by confocal microscopy, was higher in parental HeLa cells compared to GIV KO cells (**Fig 6B**; *left*; **6C**). By contrast, nuclear accumulation of Rad51 was much more pronounced in GIV KO compared to parental cells (**Fig 6B**; *right*; **6C’**). These findings indicate that NHEJ is the preferred choice for repair in cells with GIV, but HR is favored in the absence of GIV. This preference of HR over NHEJ in GIV KO cells was reversed in KO cells rescued with GIV-WT but could not be rescued by mutant GIV proteins that could not bind BRCA1 or modulate Gαi proteins (**Fig 6D**; **6E-E’**). DSBs were increased in GIV KO cells and in cells expressing either of the GIV mutants, as determined by γH2AX staining; this is consistent with the prior long amplicon PCR studies assessing the burden of mutations (**Fig 2G-H, 5G-H**).

**Figure 6.**
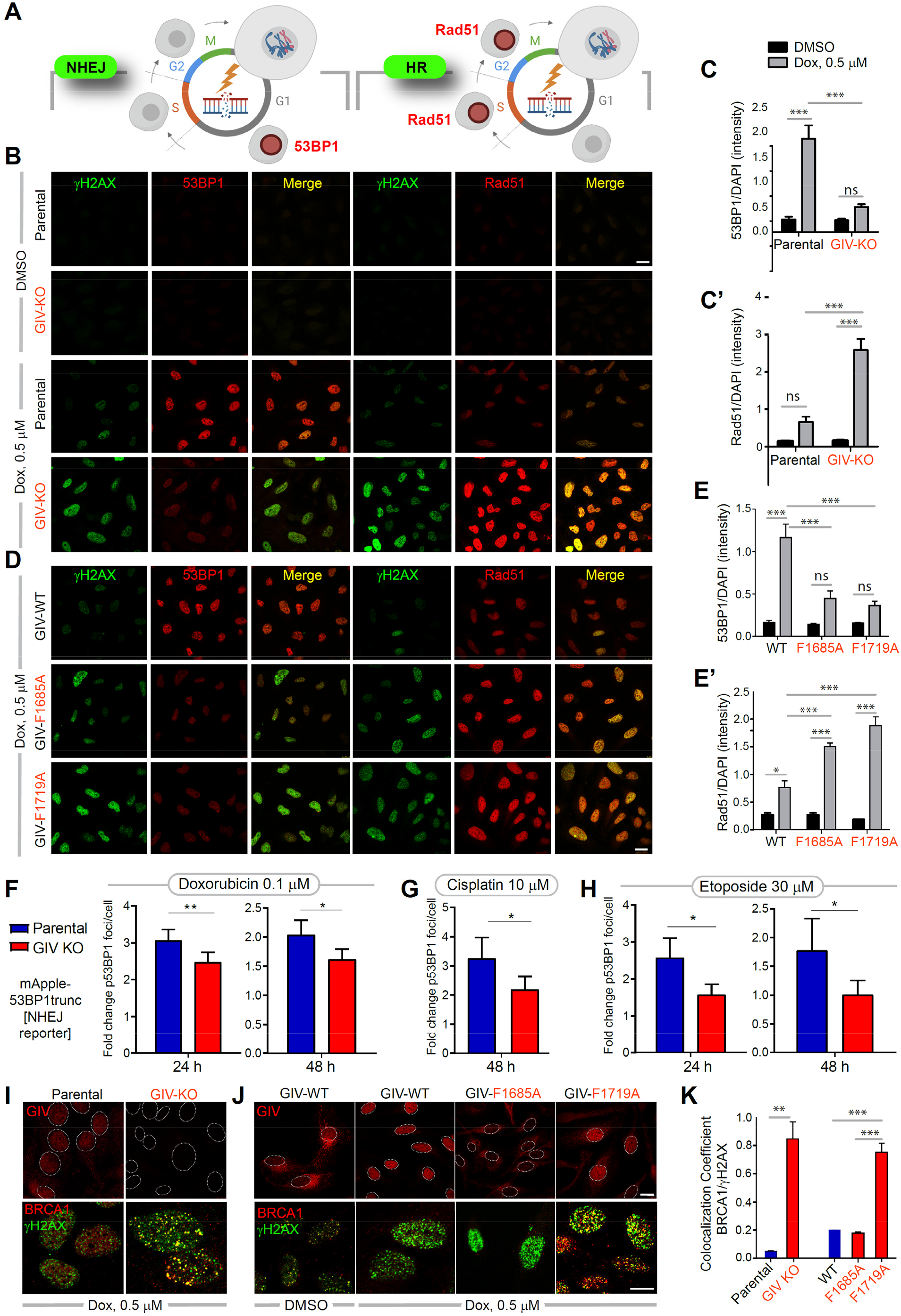
GIV inhibits HR, favors NHEJ, and inhibits localization of BRCA1 to sites of DNA damage. **A**. Schematic summarizing the two markers, 53BP1 (left) and Rad51 (right) commonly used to monitor repair pathway of choice (NHEJ vs. HR, respectively) after DNA damage. **B-E’**. Control (parental) and GIV-depleted (GIV KO) HeLa cells (B-C) or GIV-depleted HeLa cells stably expressing WT or mutant GIV constructs (D-E) were challenged with Dox or vehicle control (DMSO) prior to being fixed and co-stained for γH2AX (green) and 53BP1 (red; left) or Rad51 (red; right) and analyzed by confocal microscopy. Representative images are shown in B and D (scale bar = 15 µm). Bar graphs in C-C’ and E-E’ show the quantification of intensity of 53BP1 or Rad51 staining normalized to DAPI. Data displayed as mean ± S.E.M. and one-way ANOVA to determine significance. (*; p ≤ 0.05; **; p ≤ 0.01; ***; p ≤ 0.001; ns = not significant). **F-H**. Bar graphs display the fold change in the number of bright foci of 53BP1 in parental and GIV KO HeLa cells stably expressing mApple-53BP1 reoprter (which detects NHEJ) upon challenge with the indicated concentrations of Doxorubicin (F), Cisplatin (G) or Etoposide (H). Data displayed as mean ± S.E.M. and t-test to determine significance. (*; p ≤ 0.05; **; p ≤ 0.01). See also **Fig S7A-B** for 53BP1 reporter studies on parental and GIV KO MDA-MB-231cells. **I-K**. HeLa cell lines in B, D were treated as in B, D, and fixed and analyzed for GIV (top) and BRCA1 (bottom) localization with respect to the nuclei (demarcated with interrupted oval outlines). Representative images are shown in I-J (scale bar = 15 µm). See also **Fig S7B-C** for expanded individual panels. Bar graphs in K show Pearson’s colocalization coefficient for the degree of colocalization observed within the nucleus between BRCA1 (red) and γH2AX (green).

That GIV is required for NHEJ was further confirmed by live cell imaging using parental and GIV KO HeLa (**Fig 6F-H**) and MDA-MB-231cells (**Fig S7A-B**) stably expressing a fluorescent reporter of endogenous DSBs, a construct comprised of a truncated segment of p53BP1 fused to mApple (*29*) (see *Methods for details*). Compared to the parental cells, the fold change in bright foci/cell was significantly decreased in GIV-depleted HeLa and MDA-MB-231cells regardless of the drugs, duration and concentrations tested.

Taken together, these findings indicate that GIV and its BRCA1-binding and Gαi-modulatory functional modules are required for dictating the choice of DDR; when GIV is present and its two functional modules are intact, NHEJ is preferred over HR. It is also noteworthy that the mutational burden is increased despite the DNA damage-induced accumulation of nuclear Rad51, which suggests that HR is initiated successfully, but may not be as effective as NHEJ. The latter offers an ideal balance of flexibility and accuracy in the setting of widespread DSBs with diverse end structures (*15*).

### GIV translocates to the nucleus after DNA damage, inhibits colocalization of BRCA1 with DSBs

Because BRCA1 is a nucleocytoplasmic shuttling protein (*57*) and it is nuclear BRCA1 that augments DNA repair (*58*) and cell-cycle checkpoints (*59*), we asked if suppressed HR in cells with GIV, or those with functionally intact modules in GIV stemmed from mis-localization of BRCA1. We determined the localization of GIV and BRCA1 by confocal immunofluorescence and found that Dox challenge was associated with nuclear localization of GIV (see Parental cells; **Fig 6I**, *top-left*). Compared to parental control cells, nuclear localization of BRCA1 was more prominent in GIV KO cells (**Figure S7C**), where BRCA1 colocalized with γH2AX (see **Fig 6I**, *bottom; see* **Fig 6K** for colocalization index), indicating that nuclear localization of BRCA1 to sites of DSBs may be suppressed by GIV.

To discern which functional module of GIV may be important for nuclear localization of GIV and/or suppression of nuclear localization of BRCA1, we carried out similar assays in stable cell lines expressing GIV-WT or mutant. DNA damage dependent shuttling of GIV to the nucleus was observed in the case of GIV-WT and GIV-F1719A, but not GIV-F1685A (see **Fig 6J**, *top*), indicating that GIV’s ability to shuttle into the nucleus after DNA damage does not depend on its interaction with BRCA1, but requires a functionally intact Gαi-modulatory function. We observed prominent nuclear localization of BRCA1 only in the GIV-F1719A mutant line (**Figure S7D**), where it colocalized with γH2AX (see **Fig 6J**, *bottom*). Colocalization coefficient of BRCA1 with γH2AX across all cell lines (**Fig 6K; S7C**) showed that colocalization was greatest in the absence of GIV (GIV KO cells) or when the GIV•BRCA1 interaction is impaired (F1719A), indicating that the GIV•BRCA1 interaction is required for the observed inhibitory effect of GIV on nuclear localization of BRCA1.

Taken together, these findings demonstrate that GIV, like BRCA1, is a nucleocytoplasmic shuttling protein; shuttling is independent of its BRCA1-binding function but depends on its Gαi-modulatory function. The GIV•BRCA1 interaction appears to be primarily responsible for sequestering BRCA1 away from DSBs. Localization of BRCA1 at sites of DSBs is not only impaired in the case of GIV-WT expressing cells, in which GIV shuttles into the nucleus upon DNA damage, but also impaired in GIV-F1685A mutant cells (**Fig 6H; S7D**), in which GIV fails to localize to the nucleus. This indicates that the inhibitory GIV•BRCA1 interaction may occur in the nucleus as well as in the cytoplasm, and is in keeping with our BioID studies revealing BRCA1 as a candidate interactor of GIV in both nuclear and cytosolic compartments (**Fig 1C**).

### GIV’s Gαi-modulatory function activates Akt, BRCA1-binding function triggers S-phase checkpoint

Because the choice of DNA damage repair pathway is finetuned by a network of kinases (e.g., ATM, ATR, Akt, etc.) and the signaling cascades they initiate (*60, 61*), we asked how GIV and its functional modules may impact these pathways. More specifically, we focused on two key readouts rationalized by our observations: (i) Akt phosphorylation, because GIV is a *bona fide* enhancer of Akt phosphorylation (*62, 63*) and does so via its Gαi-modulatory function (*20*), and because this pathway is known to impact the choice of repair [reviewed in (*61*)]; (ii) Phosphorylation of the cohesion protein, Structural Maintenance of Chromosome-1 (pSer-957 SMC1) (*64*), a readout of S-phase checkpoint, because this checkpoint was impaired in GIV KO cells (**Fig 2D**). We found that the depletion of GIV significantly reduced phosphorylation of both readouts (**Fig 7A-C**), indicating that GIV is required for the phosphoactivation of both the Akt and the ATM→pSMC1 axes. Similar studies on HeLa cell lines stably expressing GIV-WT or mutants showed that Akt phosphrylation was impaired in both GIV-F1685A and GIV-F1719A mutants (**Fig 7D, 7E**), albeit more significantly impaired in the latter, but phosphoSMC1 was specifically impaired in cells expressing the GIV-F1719A mutant (**Fig 7D, 7F**). These findings show that the BRCA1-binding function of GIV is critical for initiation of Akt signaling upon DNA damage, as well as for the activation of the ATM→pSMC1 pathway for S-phase checkpoint signaling. The Gαi-modulatory function of GIV, however, was specifically responsible for enhancing Akt signals after DNA damage.

**Figure 7.**
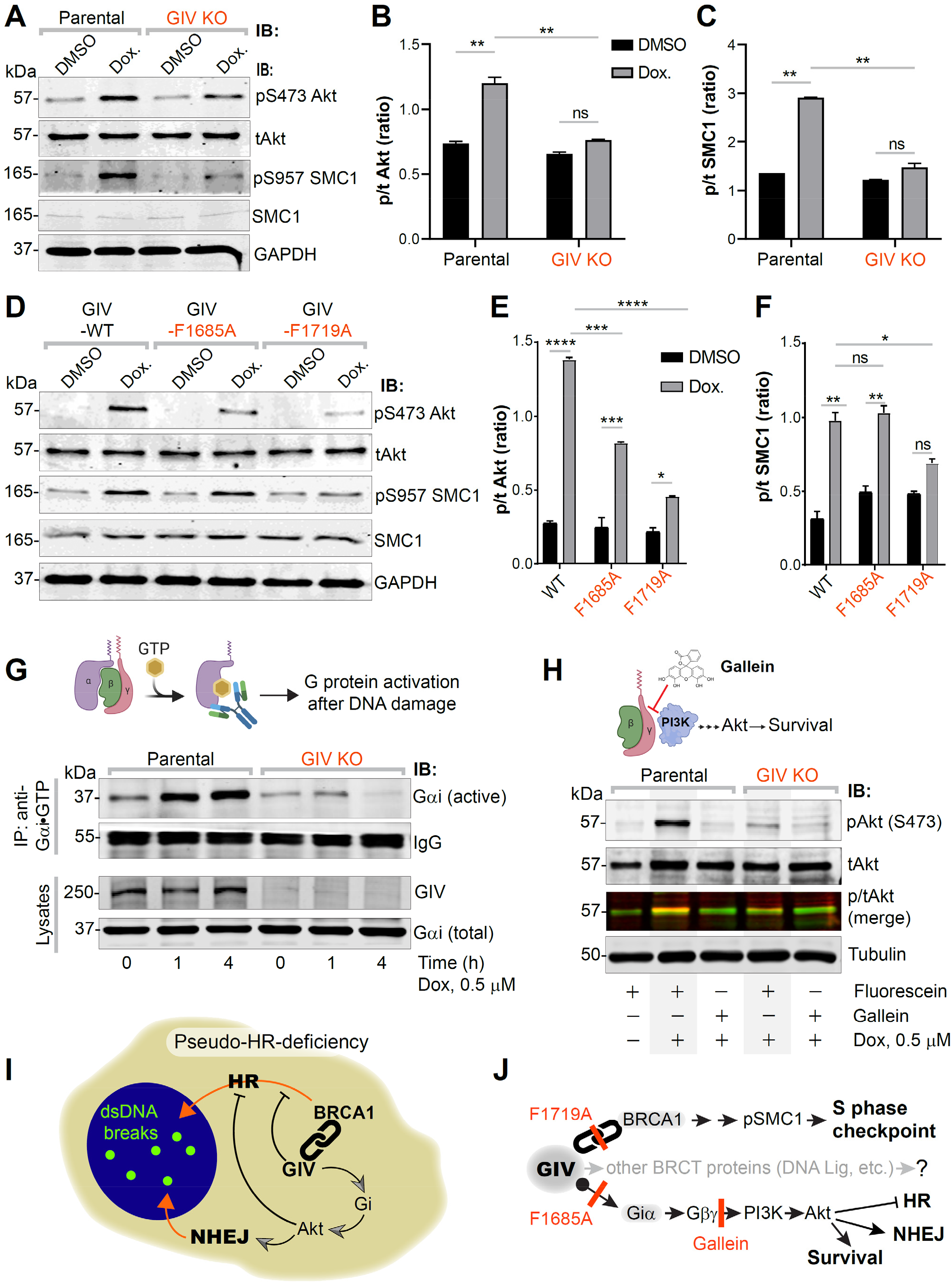
Activation of Gi by GIV is required for Akt enhancement during DDR, contributes to pseudo-HR-deficiency. **A-F**. Control (parental) and GIV-depleted (GIV KO) HeLa cells (A-C) or GIV-depleted cells stably expressing WT or mutant GIV constructs (D-F) were challenged with Dox or vehicle control (DMSO) as indicated prior to lysis. Equal aliquots of lysates were analyzed for total (t) and phosphorylated (p) Akt and SMC1 proteins and GAPDH (loading control) by quantitative immunoblotting using LiCOR Odyssey. Representative immunoblots are shown in A and D, and quantification of phospho(p)/total(t) proteins is displayed as bar graphs in B, C, E, F. **G**. Schematic on top shows the assay used for assessing the extent of Gαi-activation using conformation-sensitive antibodies that selectively bind the GTP-bound (active) conformation of Gαi protein. Immunoblots below show the active Gαi immunoprecipitated (top; IP) from lysates (bottom) of HeLa cells treated with Dox. for the indicated time points. **H**. GIV-depleted (GIV KO) and control (Parental) HeLa cells were stimulated (+) or not (-) with Dox. as indicated, in the presence of either Gallein or its inactive isomer, Fluorescein. Equal aliquots of lysates were immunoblotted for pAkt and tAkt as in panel A. I-J. Summary of findings showing how GIV skews the choice of repair pathway from HR to NHEJ, partly via sequestration of BRCA1 away from the sites of dsDNA breaks and in part via the enhancement of Akt via the Gi→’free’ GβƔ→Class I PI3K pathway. The tools (mutants and chemical inhibitors) used in this work are highlighted in red.

We asked if the previously delineated Gi → ‘free’ Gbg release → Class 1 PI3K signaling axis triggered by GIV’s Gαi-modulatory function may be essential (*20*). To this end, we first assessed the extent of activation of Gαi in cells after DNA damage by using a conformation-sensing antibody that specifically recognizes GTP-bound (active) conformation of Gαi1-3 (**Fig 7G**; *top*) (*65*), and more importantly, recognizes GIV-dependent G protein activation in cells (*48*). We found that DNA damage was associated with activation of Gαi in parental cells, but that such activation was virtually lost in GIV KO cells (**Fig 7G**; *bottom*). To dissect if Akt activation is mediated *via* the ‘free’ Gβγ→Class 1 PI3K signaling axis, we used the commonly used small molecule Gβγ inhibitor, Gallein (**Fig 7H**; *top*), and it’s an inactive isomer, Fluorescein (negative control) (*66*). We found that Gallein, but not Fluorescein inhibited DNA damage-induced Akt phosphorylation in parental control cells, reducing it to the levels observed in GIV KO cells (**Fig 7H**; *bottom*). These findings indicate that Akt signaling induced after DNA damage, occurs in part *via* GIV-dependent Gi activation.

Taken together, our findings support the following working model for how GIV dictates the choice of repair pathway after DNA damage, favoring NHEJ over HR (**Fig 7I**). Using a set of single point mutants and chemical inhibitors of G protein signaling, we charted the mechanisms that allow GIV to accomplish such a goal *via* two parallel pathways (see **Fig 7J**). One pathway is mediated by GIV’s ability to bind and sequester BRCA1 in the cytoplasmic pool, and thereby reduce its ability to localize to DSBs, suppress HR, and activate S phase checkpoint cascades. Another is GIV’s ability to bind and activate Gi and enhance Akt signaling, which further skews the choice of repair pathway towards NHEJ, while actively suppressing HR.

## Conclusion

Cellular decision-making in response to any stressful insult, is mediated by a web of spatiotemporally segregated events within the intracellular signaling networks, often requiring crosstalk between unlikely pathways. The major discovery we report here is such an unexpected crosstalk that is orchestrated *via* a versatile multi-modular signal transducer, GIV/Girdin. There are three notable takeaways from this study.

First, this work ushers a new player in DNA repair. Although GIV entered the field of cancer biology more than a decade ago, and quickly came to be known as a pro-oncogenic protein that coupled G protein signaling with unlikely pathways [reviewed in (*24*)], its role inside the nucleus remained unknown. Although predicted to have nuclear localization signals (NLS; **Supplementary Table 2**), how GIV shuttles into the nucleus remains unresolved. Regardless, what emerged using specific single-point mutants is that GIV inhibits HR by sequestering BRCA1, suppressing its localization to DSBs. Although how GIV binds BRCA1 was studied at greater depth (expanded below), how exactly GIV may inhibit the shuttling/localization of BRCA1 remains unresolved. Because nuclear import of BRCA1 and its retention requires BARD1 (*67*), whereas nuclear export requires p53 (*68*), GIV may either inhibit the BARD1•BRCA1 interplay or augment the actions of p53.

Second, one of the most unexpected observations was that GIV uses the same short linear motif (SLIM) located within its C-terminus to bind the C-terminal tandem-BRCT modules of BRCA1 in both canonical (phospho-dependent) and non-canonical (phospho-independent) modes. Although both modes of BRCT-binding has been recognized in other instances (*51*), the versatility of dual-mode binding *via* the same motif is unprecedented. Given this degree of versatility of the BRCT-binding SLIM in GIV, and the additional BRCT interactors we found here (to DNA Lig IV and BARD1), it is more likely than not that this SLIM binds other players within the DDR pathways. Which DDR proteins bind, and which do not may be dictated by the residues flanking the SLIM, as shown in other instances (*69*). These findings are in keeping with the fact that GIV-CT is an intrinsically disordered protein (IDP) (*47, 70*) comprised of distinct SLIMs, of which the BRCT-binding motif described here is an example (see **Fig S3A**). SLIMs enable GIV to couple G protein signaling to a myriad of molecular sensors, of both the outside of the cell (i.e., receptors; [reviewed in (*71*)]) or its interior (*72*). Because IDPs that fold/unfold on demand expose/hide SLIMs, which in turn imparts plasticity to protein-protein interaction networks during signal transduction (*73*), GIV may do something similar in couple G protein signaling to DDR. By scaffolding G proteins to BRCT-modules in BRCA1 (and presumably other DDR proteins) GIV may serve as a point of convergence for coordinating signaling events and generating pathway crosstalk upon DNA damage.

Third, this work provides a direct mechanistic link between DDR and trimeric G proteins; the latter is one of the major pervasive signaling hubs in eukaryotic cells that was notably absent from the field of DNA repair. Although multiple peripheral components within the GPCR/G-protein signaling system has been found to indirectly influence DNA damage and/or repair (*23*), who/what might activate G proteins on endomembranes was unknown. We demonstrated that trimeric Gi proteins are activated upon DNA damage, and that such activation requires GIV’s Gαi-modulatory motif. That the GIV→Gαi pathway activates Akt signaling helps explain the hitherto elusive origin of Akt signaling during DDR (*61*). That GIV favors NHEJ over HR and activates Akt signaling during DDR is in keeping with the previously described role of Akt signaling in dictating the choice of repair pathway (*61*).

In closing, damage to the genome can have catastrophic consequences, including cytotoxicity, accelerated aging, and predisposition to cancers. Our findings, which revealed a hitherto unknown link between a major hub in DNA repair (i.e., BRCA1) and a signaling hub of paramount importance in just about all aspects of modern medicine (trimeric G proteins) opens new avenues for development of novel therapeutic strategies.

## AUTHOR CONTRIBUTIONS

A. A. A. and P.G designed, executed and analyzed most of the experiments in this work. N.S. carried out all work related to the generation of constructs used in this work. N.S, P.C., and N.R conducted the protein chemistry and biochemical analyses of the protein-protein interactions. A.C. and A.A.A. designed, executed, and analyzed the DNA mutation load assays in consultation with S.D and supervised by T.K.H. J.E designed, executed, and analyzed the BioID studies. S.R. carried out the structural modeling and analyses with supervision from P.G. K.E.L and G.D.L designed, executed, and analyzed the 53BP1 reporter studies in live cells. A. A. A., A.C and S.R wrote methods. A.A.A and P.G conceived the project, wrote and edited the manuscript.

## Corresponding author

Correspondence to Pradipta Ghosh

## Data availability

All data are included in the manuscript or deposited as extended data.

