## Supplementary Online Materials for "Regulation of DNA damage response by trimeric G-protein Signaling"

### INVENTORY OF SUPPLEMENTARY MATERIALS


- TRANSPARENT METHODS
- SUPPLEMENTARY TABLES (2)
- SUPPLEMENTARY FIGURES AND LEGENDS (8)
- LEGENDS FOR EXTENDED DATA (2)

### TRANSPARENT METHODS

- **Key Resource Table**
- **Contact for Reagent and Resource Sharing**
- **Experimental Model and Subject Details**
  - Cell Lines (HeLa, Cos7, HEK, Hs578T)
- **Method Details**
  - *Cell lines and culture methods*
  - *Plasmid constructs and mutagenesis*
  - *Transfection, generation of stable cell lines and cell lysis*
  - *Biotin Proximity Labeling*
  - *In Gel Digest*
  - *LC-MS analysis*
  - *Gene Ontology Analysis*
  - *GIV CRISPR/Cas9 Gene Editing and Validation*
  - *Protein expression and purification*
  - *In Vitro GST-Pulldown and In-cellulo Co-immunoprecipitation (CoIP) Assays*
  - *Quantitative immunoblotting*
  - *Immunofluorescence and Confocal Microscopy, Image analysis*
  - *MTT assays*
  - *Cell cycle and apoptosis analyses*
  - *Long Amplicon PCR*
  - *Using Human Cell Map for the identification of potential interactors and subcellular compartment annotation of GIV*
  - *G protein activity assay*
  - *Homology modeling*
  - *Image Processing*
- **Quantification and Statistical Analysis**
  - Statistical Analysis
  - Replications
- **Data and Software Availability**

### Key resource table

| REAGENT or RESOURCE | SOURCE | IDENTIFIER |
|---|---|---|
| **ANTIBODIES** | | |
| **Rabbit polyclonal anti-GIV (Girdin) (T-13)** | Santa Cruz Biotechnology | sc-133371 |
| **Rabbit polyclonal anti-GIV (Girdin) (CC-Ab)** | Millipore Sigma | ABT80 |
| **Rabbit polyclonal anti-GIV (CT-Ab)** | *This paper* | N/A |
| **Mouse monoclonal anti-Myc** | Cell Signaling Technology | 2276S |
| **Mouse monoclonal anti-GAPDH** | Santa Cruz Biotechnology | sc-365062 |
| **Rabbit polyclonal anti-BRCA1** | Santa Cruz Biotechnology | sc-642 |
| **Mouse monoclonal anti-α-tubulin** | Santa Cruz Biotechnology | sc-5286 |
| **Mouse monoclonal anti-FLAG** | Millipore Sigma | MAB3118 |
| **Mouse monoclonal anti-Myc** | Cell Signaling Technology | 2276S |
| **Mouse monoclonal anti-GST** | GenScript | A00865 |
| **Rabbit polyclonal anti-BACH1** | Proteintech | 14018-1-AP |
| **Rabbit polyclonal anti-pan Gβ** | Proteintech | sc-378 |
| **Rabbit polyclonal anti-GFP(CtIP)** | Santa Cruz Biotechnology | sc-9996 |
| **Mouse monoclonal anti-p53** | Santa Cruz Biotechnology | sc-99 |
| **Mouse monoclonal anti-p-YH2A.X** | Santa Cruz Biotechnology | sc-517348 |
| **Rabbit polyclonal anti-53BP1** | Cell Signaling Technology | 4937S |
| **Rabbit polyclonal anti-RAD51** | Proteintech | 14961-1-AP |
| **Rabbit monoclonal anti-pS473 AKT** | Cell Signaling Technology | D9E |
| **Mouse monoclonal anti-total AKT** | Cell Signaling Technology | 40D4 |
| **Mouse monoclonal anti-pS957 SMC** | Cell Signaling Technology | 5D11G5 |
| **Mouse monoclonal anti-SMC** | Cell Signaling Technology | 8E6 |
| **Rabbit polyclonal anti-Gαi3 (C-10)** | Santa Cruz Biotechnology | N/A |
| **Rabbit polyclonal anti-Gαi (total)** | Santa Cruz Biotechnology | Sc-389 |
| **Mouse monoclonal anti- Gαi-GTP** | *Graeme Milligan* (*65*) | 26901 |
| **Goat anti-Rabbit IgG, Alexa Fluor 594 conjugated** | ThermoFisher Scientific | A11072 |
| **Goat anti-Mouse IgG, Alexa Fluor 488 conjugated** | ThermoFisher Scientific | A11017 |
| **IRDye 800CW Goat anti-Mouse IgG Secondary (1:10,000)** | LI-COR Biosciences | 926-32210 |
| **IRDye 680RD Goat anti-Rabbit IgG Secondary (1:10,000)** | LI-COR Biosciences | 926-68071 |
| **BIOLOGICAL SAMPLES AND CELL LINES** | | |
| **CELL LINE** | Source | Cat# |
| **HeLa parental** | ATCC | ATCC® CCL-2 |
| **HeLa GIV KO (CRISPR Cas9)** | *This paper* | n/a |
| **HeLa GIV WT** | *This paper* | n/a |
| **HeLa GIV F1685A** | *This paper* | n/a |
| **HeLa GIV F1719A** | *This paper* | n/a |
| **HEK293T** | ATCC | ATCC® CRL-11268 |
| **COS7** | ATCC | ATCC® CRL-1651 |
| **DLD1** | ATCC | ATCC® HTB-126 |
| **DLD1 parental and GIV KO (CRISPR Cas9) lines** | n/a | Published in (*28*) |

| | | |
|---|---|---|
| **MDA-MB-231** | ATCC | ATCC® HTB-26 |
| **MDA-MB-231 parental and GIV KO (CRISPR Cas9) lines** | *This paper* | n/a |
| **CHEMICALS, KITS, RECOMBINANT PROTEINS** | | |
| **Biotin** | Sigma-Aldrich | B4639-500MG |
| **DAPI (4',6-Diamidino-2-Phenylindole, Dilactate)** | Thermo Fisher Scientific | D3571 |
| **G418** | Cellgro | A-1720 |
| **Paraformaldehyde 16%** | Electron Microscopy Biosciences | 15710 |
| **DIG RNA Labeling Mix** | Roche | 11277073910 |
| **T7 RNA polymerase** | Promega | P2075 |
| **Doxorubicin** | Sigma Aldrich | D1515-10MG |
| **Gallein** | TCI Chemicals | 2103-64-2 |
| **Fluorescein** | TCI Chemicals | 2321-07-5 |
| **MTT** | Millipore Sigma | 475989-1GM |
| **Cisplatin** | EMD Millipore | 232120-50MG |
| **Etoposide** | Sigma-Aldrich | E1383-25MG |
| **Guava Cell Cycle Reagent** | Millipore Sigma | 4700-0160 |
| **Dead Cell Apoptosis Kit with Annexin V Alexa Fluor™ 488 & Propidium Iodide (PI)** | ThermoFisher Scientific | V13241 |
| **DAPI (4',6-Diamidino-2-Phenylindole, Dilactate)** | Thermo Fisher Scientific | D3571 |
| **Streptavidin, Alexa Fluor® 680 conjugate** | ThermoFisher Scientific | S21378 |
| **Streptavidin, Alexa Fluor® 594 conjugate** | ThermoFisher Scientific | S11227 |
| **HisPur[ä] Cobalt Resin** | Thermo Scientific | 89964 |
| **Glutathione Sepharose[â] 4B** | Sigma-Aldrich | GE17-0756-04 |
| **Streptavidin Magnetic Beads** | ThermoFisher Scientific | 88816 |
| **Protein A Agarose** | ThermoFisher | 15918014 |
| **Protein G Agarose** | ThermoFisher | 20398 |
| **Protease inhibitor cocktail** | Roche | 11 873 580 001 |
| **Tyr phosphatase inhibitor cocktail** | Sigma-Aldrich | P5726 |
| **Ser/Thr phosphatase inhibitor cocktail** | Sigma-Aldrich | P0044 |
| **PVDF Transfer Membrane, 0.45mM** | Thermo Scientific | 88518 |
| **PLASMIDS AND CONSTRUCTS** | | |
| **Girdin CRISPR/Cas9 KO Plasmid (h2)** | *Santa Cruz Biotechnology (SCBT) Inc.* | Sc-402236-KO-2 |
| **GST-BRCA1-CT-WT (BRCT) [pGEX-4T-human BRCA1 BRCT domain (residues 1599-1863)]** | *Zhou Songyang* (*74*) | N/A |
| **GST-BRCA1-CT-M1775R (BRCT) [pGEX-4T-human BRCA1 BRCT domain (residues 1599-1863)]** | *This paper* | N/A |
| **GST-BRCA1-NT(RING) [pGEX-4T-human BRCA1 RING domain (residues 1-661)]** | *Zhou Songyang* (*74*) | N/A |
| **GST-MDC1 [pGEX-4T-human MDC1 (residues 2727-3089)]** | *Zhou Songyang* (*74*) | N/A |
| **GST-DNA ligase IV [pGEX-4T-human DNA ligase IV (residues 618-911)]** | *Zhou Songyang* (*74*) | N/A |
| **GST-BARD1 [pGEX-4T-human BARD1 (residues 554 to 777)** | *Zhou Songyang* (*74*) | N/A |
| **BACH1/FANCJ myc tagged pcDNA3 vector** | *David Livingston's group (75)* | N/A |

| | | |
|---|---|---|
| **GFP-tagged CtIP** | *David Livingston's group (76)* | N/A |
| **HA-tagged BRCA1-WT** | *David Livingston's group (77)* | N/A |
| **pcDNA3(*BssH*II)-HA-3XFLAG-BRCA1 WT** | *Kristoffer Valerie's group (50)* | N/A |
| **pcDNA3(*BssH*II)-HA-3XFLAG-BRCA1 K1702M** | *Kristoffer Valerie's group* Dever, 2011 #7} | N/A |
| **CMV14-p3X FLAG-GIV WT (full length)** | *(78)* | N/A |
| **CMV14-p3X FLAG-GIV-F1685A (full length)** | *(78)* | N/A |
| **CMV14-p3X FLAG-GIV-F1719A (full length)** | *This paper* | N/A |
| **CMV14-p3X FLAG-GIV-S1716D (full length)** | *This paper* | N/A |
| **CMV14-p3X FLAG-GIV-S1716D/F1719A (full length)** | *This paper* | N/A |
| **pET-28b-GIV-CT-WT (aa 1623-1870)** | *(78)* | N/A |
| **pET-28b-GIV-CT-WT (aa 1660-1870)** | *(78)* | N/A |
| **pET-28b-GIV-CT-WT (aa 1790-1870)** | *(78)* | N/A |
| **pGEX-4T-GIV-CT-WT (a.a. 1623-1870)** | *(78)* | N/A |
| **pGEX-4T-GIV-CT-F1685A (a.a. 1623-1870)** | *(78)* | N/A |
| **pGEX-4T-GIV-CT-S1716A (a.a. 1623-1870)** | *This paper* | N/A |
| **pGEX-4T-GIV-CT-S1716D (a.a. 1623-1870)** | *This paper* | N/A |
| **pET-28b-GIV-CT-F1685A (aa 1660-1870)** | *(78)* | N/A |
| **pET-28b-GIV-CT-S1716D (aa 1660-1870)** | *This paper* | N/A |
| **pET-28b-GIV-CT-S1716A (aa 1660-1870)** | *This paper* | N/A |
| **Lentiviral vector expressing a truncated version of p53BP1 fused to mApple** | *Addgene* | *69531* |
| **Prolong Glass** | Thermo Fisher Scientific | P36980 |
| **Paraformaldehyde 16%** | Electron Microscopy Biosciences | 15710 |
| **SOFTWARE** | | |
| **ImageJ** | National Institute of Health | https://imagej.net/Welcome |
| **DAVID 6.8** | DAVID Bioinformatics Resources | https://david.ncifcrf.gov/home.jsp |
| **FlowJo** | FlowJo, LLC | https://www.flowjo.com |
| **Prism** | GraphPad | https://www.graphpad.com/scientific-software/prism/ |
| **LAS-X** | Leica | www.leica-microsystems.com/products/microscope-software/p/leica-las-x-ls |
| **Molsoft** | Molsoft, LLC | https://www.molsoft.com/index.html |
| **Pymol** | Pymol.org | https://pymol.org/2/ |
| **Illustrator** | Adobe | https://www.adobe.com/products/illustrator.html |
| **ImageStudio Lite** | LI-COR | https://www.licor.com/bio/image-studio-lite/ |
| **MATLAB** | MathWorks | https://www.mathworks.com/products/matlab.html |

### Contact for Reagent and Resource Sharing

Pradipta Ghosh

**Method details:**

*Cell lines and culture methods*

HeLa, Cos7, HEK, Hs578T, DLD-1, and MDA-MB-231 cells were grown at 37°C in their suitable media, according to their supplier instructions, supplemented with 10% FBS, 100 U/ml penicillin, 100 µg/ml streptomycin, 1% L-glutamine, and 5% CO2.

*Plasmid constructs and mutagenesis*

For mammalian expression, a well-characterized and extensively validated C-terminal FLAG-tagged construct (Ghosh et al., 2010) was used. It was originally generated by cloning human GIV (NCBI RefSeq Accession: Q3V6T2) into p3XFLAG-CMV-14 between NotI and BamHI. All subsequent site-directed mutagenesis (GIV-Flag full-length F1685A, F1719A, S1716A (SA), and S1716D (SD) were carried out on this template using Quick Change as per the manufacturer's protocol. For BRCA1 constructs, the sources are listed in the "Table of Key Resources". The target mutants for GIV and BRCA1 were confirmed by sequencing. The GST-BRCA1-WT, GST-BRCA1-CT (BRCT), GST-BRCA1-NT, GST-BRCA1-K1702M, GST-BRCA1-M1775R, GST-MDC1, GST-DNA ligase IV, GST-BARD1, GST-GIV CT, GST-GIV WT, GST-GIV F1685A, GST-GIV F1719A, GST-GIV-CT-S1716A, GST-GIV-CT- S1716D, His GIV-CT, and GST-Gαi3 fusion proteins were used for in vitro protein-protein interaction.

*Transfection, generation of stable cell lines and cell lysis*

Transfection was carried out using Genejuice (Novagen) for DNA plasmids following the manufacturers' protocols. Hela cell lines stably expressing GIV constructs (WT, F1685A, and F1719A) were selected after transfection in the presence of 800 mg/ml G418 for 6 weeks. The resultant multiclonal pool was subsequently maintained in the presence of 500 mg/ml G418. GIV expression was verified independently by immunoblotting using anti-GIV antibody. Whole-cell lysates were prepared after washing cells with cold PBS prior to resuspending and boiling them in sample buffer. Lysates used as a source of proteins in immunoprecipitation or pulldown assays were prepared by resuspending cells in Tx-100 lysis buffer [20 mM HEPES, pH 7.2, 5 mM Mg-acetate, 125 mM K-acetate, 0.4% Triton X-100, 1 mM DTT, supplemented with sodium orthovanadate (500 mM), phosphatase (Sigma) and protease (Roche) inhibitor cocktails], after which they were passed through a 28G needle at 4˚C, and cleared (10,000 x g for 10 min) before use in subsequent experiments.

*Biotin Proximity Labeling*

BioID was performed as previously described (*27*). Briefly, HEK293T were plated 24 hrs prior to transfection with mycBirA-tagged GIV construct. Thirty hours post transfection, cells were treated with 50 µM biotin (dissolved in culture media) for 16 hrs. Cells were then rinsed two times with PBS and lysed by resuspending in lysis buffer (50 mM Tris, pH 7.4, 500 mM NaCl, 0.4% SDS, 1 mM dithiothreitol, 2% Triton X-100, and 1× Complete protease inhibitor) and sonication in a bath sonicator. Cell lysates were then cleared by centrifugation at 20,000 X g for 20 mins and supernatant was then collected and incubated with streptavidin magnetic beads overnight at 4°C.

After incubation, beads were washed twice with 2% SDS, once with wash buffer 1 (0.1% deoxycholate, 1% Triton X-100, 500 mM NaCl, 1 mM EDTA, and 50 mM HEPES, pH 7.5), followed with once wash using wash buffer 2 (250 mM LiCl, 0.5% NP-40, 0.5% deoxycholate, 1 mM EDTA, and 10 mM Tris, pH 8.0), and once with 50 mM Tris pH 8.0. Biotinylated complexes were then eluted using sample buffer containing excess biotin and heating at 100°C. Prior to mass spectrometry identification, eluted samples were run on SDS-PAGE and proteins were extracted by in gel digest.

*In Gel Digest*

Protein digest and mass spectrometry was perform as previously described (*79*). Briefly, the gel slices were cut into 1mm X 1 mm cubes, destained 3 times by first washing with 100 ul of 100 mM ammonium bicarbonate for 15 minutes, followed by the addition of equal volume acetonitrile (ACN) for 15 minutes. The supernatant was collected, and samples were dried using a speedvac. Samples were then reduced by mixing with 200 µl of 100 mM ammonium bicarbonate-10 mM DTT and incubated at 56°C for 30 minutes. The liquid was removed and 200 ul of 100 mM ammonium bicarbonate-55mM iodoacetamide was added to gel pieces and incubated covered at room temperature for 20 minutes. After the removal of the supernatant and one wash with 100 mM ammonium bicarbonate for 15 minutes, equal volume of ACN was added to dehydrate the gel pieces. The solution was then removed, and samples were dried in a SpeedVac. For digestion, enough solution of ice-cold trypsin (0.01 ug/ul) in 50 mM ammonium bicarbonate was added to cover the gel pieces and set on ice for 30 min. After complete rehydration, the excess trypsin solution was removed, replaced with fresh 50 mM ammonium bicarbonate, and left overnight at 37°C. The peptides were extracted twice by the addition of 50 µl of 0.2% formic acid and 5 % ACN and vortex mixing at room temperature for 30 min. The supernatant was removed and saved. A total of 50 µl of 50% ACN-0.2% formic acid was added to the sample, and vortexed again at room temperature for 30 min. The supernatant was removed and combined with the supernatant from the first extraction. The combined extractions are analyzed directly by liquid chromatography (LC) in combination with tandem mass spectroscopy (MS/MS) using electrospray ionization.

*LC-MS analysis*

Trypsin-digested peptides were analyzed by ultra-high-pressure liquid chromatography (UPLC) coupled with tandem mass spectroscopy (LC-MS/MS) using nano-spray ionization. The nanospray ionization experiments were performed using a Orbitrap fusion Lumos hybrid mass spectrometer (Thermo) interfaced with nano-scale reversed-phase UPLC (Thermo Dionex UltiMate™ 3000 RSLC nano System) using a 25 cm, 75-micron ID glass capillary packed with 1.7-µm C18 (130) BEH™ beads (Waters corporation). Peptides were eluted from the C18 column into the mass spectrometer using a linear gradient (5–80%) of ACN (Acetonitrile) at a flow rate of 375 µl/min for 1h. The buffers used to create the ACN gradient were: Buffer A (98% $H_2O$, 2% ACN, 0.1% formic acid) and Buffer B (100% ACN, 0.1% formic acid). Mass spectrometer parameters are as follows; an MS1 survey scan using the orbitrap detector (mass range (m/z): 400-1500 (using quadrupole isolation), 120000 resolution setting, spray voltage of 2200 V, Ion transfer tube temperature of 275 C, AGC target of 400000, and maximum injection time of 50 ms) was followed by data dependent scans (top speed for most intense ions, with charge state set to only include +2-5 ions, and 5 second exclusion time, while selecting ions with minimal intensities of 50000 at in which the collision event was carried out in the high energy collision cell (HCD Collision Energy of 30%), and the fragment masses where analyzed in the ion trap mass analyzer (With ion trap scan rate of turbo, first mass m/z was 100, AGC Target 5000 and maximum injection time of 35ms). Protein identification and label free quantification was carried out using Peaks Studio 8.5 (Bioinformatics solutions Inc.)

*Gene Ontology Analysis*

Identified proteins by mass spec. anaylsis unique to plus biotin samples, but not in minus biotin samples, were analyzed using DAVID and functional annotation was grouped by molecular function and cellular component for GO analysis. Classification with p-value less than 0.05 were considered as significant.

*GIV CRISPR/Cas9 Gene Editing and Validation*

Pooled guide RNA plasmids (commercially obtained from Santa Cruz Biotechnology; Cat# sc-402236-KO-2) were used. These CRISPR/Cas9 KO plasmids consists of GFP and girdin-specific 20 nt guide RNA sequences derived from the GeCKO (v2) library and target human Girdin exons 6 and 7. Plasmids were transfected into Hela and MDA-MB-231 cells using PEI. Cells were sorted into individual wells using a cell sorter based on GFP expression. To identify cell clones harboring mutations in gene coding sequence, genomic DNA was extracted using 50 mM NaOH and boiling at 95°C for 60mins. After extraction, pH was neutralized by the addition of 10% volume 1.0 M Tris-pH 8.0. The crude genomic extract was then used in PCR reactions with primers flanking the targeted site. Amplicons were analyzed for insertions/deletions (indels) using a TBE-PAGE gel. Indel sequence was determined by cloning amplicons into a TOPO-TA cloning vector (Invitrogen) following manufacturer's protocol. DLD1 parental and GIV KO lines were generated and validated as described before (*28*).

*Protein expression and purification*

GST and His-tagged recombinant proteins were expressed in E. coli strain BL21 (DE3) (Invitrogen) and purified as described previously (*20, 21*). Briefly, bacterial cultures were induced overnight at 25˚C with 1 mM isopropylb-D-1-thio-galactopyranoside (IPTG). Pelleted bacteria from 1 L of culture were resuspended in 20 mL GST-lysis buffer [25 mM TrisHCl, pH 7.5, 20 mM NaCl, 1 mM EDTA, 20% (vol/vol) glycerol, 1% (vol/vol) Triton X-100, 2X protease inhibitor mixture (Complete EDTA-free; Roche Diagnostics)] or in 20 ml His-lysis buffer [50 mM NaH2PO4 (pH 7.4), 300 mM NaCl, 10 mM imidazole, 1% (vol/vol) Triton X-100, 2X protease inhibitor mixture (Complete EDTA-free; Roche Diagnostics)] for GST or His-fused proteins, respectively. After sonication (three cycles, with pulses lasting 30 s/cycle, and with 2 min intervals between cycles to prevent heating), lysates were centrifuged at 12,000X g at 4˚C for 20 min. Solubilized proteins were affinity purified on glutathione-Sepharose 4B beads (GE Healthcare) or HisPur Cobalt Resin (Pierce), dialyzed overnight against PBS, and stored at 80˚C.

*In Vitro GST-Pulldown and In-cellulo Co-immunoprecipitation (CoIP) Assays*

Purified GST-tagged proteins from E. coli were immobilized onto glutathione-Sepharose beads and incubated with binding buffer (50 mM Tris-HCl (pH 7.4), 100 mM NaCl, 0.4% (v:v) Nonidet P-40, 10 mM MgCl2, 5 mM

EDTA, 2 mM DTT) for 60 mins at room temperature. For the pulldown of protein-protein complexes from cell lysates, cells were first lysed in cell lysis buffer (20 mM HEPES, pH 7.2, 5 mM Mg-acetate, 125 mM K-acetate, 0.4% Triton X-100, 1 mM DTT, 500 µM sodium orthovanadate, phosphatase inhibitor cocktail (Sigma-Aldrich) and protease inhibitor cocktail (Roche)) using a 28G needle and syringe, followed by centrifugation at 10,000Xg for 10 mins. Cleared supernatant was then used in binding reaction with immobilized GST-proteins for 4 hours at 4°C. After binding, bound complexes were washed four times with 1 ml phosphate wash buffer (4.3 mM Na2HPO4, 1.4 mM KH2PO4, pH 7.4, 137 mM NaCl, 2.7 mM KCl, 0.1% (v:v) Tween 20, 10 mM MgCl2, 5 mM EDTA, 2 mM DTT, 0.5 mM sodium orthovanadate). Bound proteins were then eluted through boiling at 100°C in sample buffer.

For CoIP assays, cells lysates (as prepared above) was incubated with capture antibodies for 3 hours at 4°C, followed by the addition of Protein A or Protein G beads to capture antibody bound protein-protein complexes. Bound proteins were then eluted through boiling at 100°C in sample buffer.

*Quantitative immunoblotting*

For immunoblotting, protein samples were boiled in Laemmli sample buffer, separated by SDS-PAGE and transferred onto 0.4µm PVDF membrane (Millipore) prior to blotting. Post transfer, membranes were blocked using 5% Non-fat milk or 5% BSA dissolved in PBS. Primary antibodies were prepared in blocking buffer containing 0.1% Tween-20 and incubated with blots, rocking overnight at 4°C. After incubation, blots were incubated with secondary antibodies for one hour at room temperature, washed, and imaged using a dual-color Li-Cor Odyssey imaging system.

*Immunofluorescence and Confocal Microscopy, Image analysis*

Cells were fixed using -20°C methanol (or 4°C paraformaldehyde, PFA) for 20 to 30 mins, rinse with PBS, and then permeabilized for 1hr using blocking/permeabilization buffer (0.4% Triton X-100 and 2 mg/ml BSA dissolved in PBS). Primary antibody and secondary antibody were diluted in blocking buffer and incubated with cells for 1 hr each. Coverslips were mounted using Prolong Gold (Invitrogen) and imaged using a Leica SPE CTR4000 confocal microscope.

*MTT assay*

Cell proliferation was measured using the MTT reagent and cells cultured in 96-well plates. Parental or GIV-KO HeLa, DLD1 or MDA-MB-231 cells or HeLa GIV-KO cells stably expressing WT GIV, GIV F1685A, or GIV F1719A were cultured and treated with different concentrations of Doxorubicin (0.1, 0.25, 0.5, 1, 2, and 4 µM), cisplatin (1, 5, 10, 25, 50, and 100µM), or etoposide (1, 5, 10, 25, 50, and 100µM) using DMSO as a negative control for 24h. MDA-MB-231 and DLD-1 cell lines were treated with only dox (0.5µM) for 24h. Then the cell lines were incubated with MTT for 4 hr at 37°C. After incubation, culture media was removed and 150 µl of DMSO was added in order to solubilize the MTT formazan crystals. Optical density was determined at 590 nm using a TECAN plate reader. At least three independent experiments were performed.

*Anchorage-dependent colony formation assay*

Anchorage-dependent growth was monitored as described previously (*21, 80*). Briefly, anchorage-dependent growth was monitored on solid (plastic) surface. Approximately 2,000 parental or GIV-KO HeLa cells or GIV-KO cells stably expressing WT GIV, GIV F1685A, or GIV F1719A were plated in 6-well plates and incubated in 5% CO2 at 37°C for ~2 weeks in 2% FBS growth media in the presence of 10 nM Dox. After every three days, media were changed with fresh media containing 10 nM Dox. Colonies were then stained with 0.005% crystal violet for 1 hr. Entire plate surface area was scored for colonies and each treatment was done in triplicate and repeated thrice.

*Cell cycle and apoptosis analyses*

Cell cycle analysis and apoptotic cell quantification was performed using the Guava cell cycle reagent (Millipore Sigma) or the annexin V/propidium iodide (PI) staining kit (Thermo Fisher Scientific), respectively, according to the manufacturer's instructions. Cells were quantified on a BD LSR II flow cytometer and analyzed using FlowJo software (FlowJo, Ashland, OR, USA).

*Long Amplicon PCR*

Genomic DNA extraction was performed using the genomic-tip 20/G kit (Qiagen, Cat no. 10223, with corresponding buffer sets) per the manufacturer's directions. This kit has the advantage of minimizing DNA oxidation during the isolation steps, and thus it can be used reliably for isolation of high molecular weight DNA with excellent template integrity to detect endogenous DNA damage using LA-qPCR. After precise quantitation of the DNA by Pico Green (Invitrogen Cat no. P7589) in a 96-well black-bottomed plate, the genomic DNA (500 ng) was digested with the E. coli enzymes Fpg and Nei (New England Biolabs) in reaction volume of 50 uL using Buffer 1 from NEB (with 1 mM MgCl2) as the common buffer to induce strand breaks at the sites of the unrepaired oxidized base lesion. Gene-specific LA-qPCR analyses for measuring DNA damage were performed using Long Amp Taq DNA polymerase (New England Biolabs, Cat no MO323S). The numbers of cycles and DNA concentrations were standardized in each case before the actual reaction, so that the PCR remains within the linear range of amplification. The final PCR condition was optimized at 94 °C for 30 s (94 °C for 30 s, 55–60 °C for 30 s depending on the oligo annealing temperature, 65 °C for 10 min) for 25 cycles and 65 °C for 10 min. 25 ng of DNA template was used in each case, and the LA-qPCR was set for all the genes under study from the same stock of Fpg/Nei-treated diluted genomic DNA samples to avoid variations in PCR amplification due to sample preparation. Since amplification of a small region would be independent of DNA damage, a small DNA fragment for each gene was also amplified to normalize the amplification of large fragments. The PCR conditions were 94 °C for 30 s (94 °C for 30 s, 54 -58°C for 20 s, and 68 °C for 30 s) for 25 cycles and 68 °C for 5 min. 25 ng of template from the same Fpg/Nei digested DNA aliquot was used for short PCR using green mix (NEB). The amplified products were then visualized on gels and quantitated with an ImageJ automated digitizing system (National Institutes of Health) based on three independent replicate PCRs. The extent of damage was calculated.

*Stable cell lines with p53BP1 fluorescent reporter*

We obtained a lentiviral vector expressing a truncated version of p53BP1 fused to mApple from Dr. Ralph Weissleder, Massachusetts General Hospital, USA (Addgene #69531) (*29*). To establish MDA-MB-231 and HeLa parental and GIV knockout cells stably expressing the reporter, we produced recombinant lentiviruses and transduced each cell line as described previously (*30*). We selected batch populations of transduced cells with 5 μg/ml puromycin. After selection, we maintained cells in DMEM (#10313, Gibco, Thermo Fisher, Grand Island, NY, USA) with 10% FBS (HyClone, ThermoScientific, Waltham, MA, USA), 1% GlutaMAX (#35050, Gibco), and 1% Penicillin-Streptomycin (P/S, #15140, Gibco) and no added puromycin.

*Image acquisition and analysis*

We seeded cells in 6- or 96-well glass bottom plates (P-96-1.5H-N or P06-1.5H-N, Cellvis, Mountain View, CA, USA) at densities of 1.25 x 105 or 5 x 103 cells/well, respectively, in imaging base medium (FluoroBrite DMEM media (A1896701, ThermoFisher Scientific, Waltham, MA USA), 1% GlutaMax, 1% PenStrep, 1% sodium pyruvate, and 10% FBS (HyClone)). One day later, we acquired baseline pre-treatment images and then changed to fresh imaging base medium containing types and concentrations of chemotherapy drugs indicated in figure legends. We repeated imaging after one day of treatment and in selected experiments imaged again after two days. We performed imaging studies with an EVOS M7000 Imaging System (ThermoFisher), 40X objective, and the RFP cube for the instrument. During imaging, we used incubator conditions of 37 ᵒC, 5% CO2, and 80% humidity. We randomly acquired Z-stack images (7-9 planes at ~ 0.8 μm intervals) from 6-8 fields per condition, which typically encompassed > 100 cells each. To accurately quantify the number of bright P53BP1 foci per cell nucleus, we developed custom MATLAB image processing and analysis software. To process the raw images, we first smoothed each plane with an edge-preserving Gaussian bilateral filter and then calculated a maximum intensity projection (MIP) from these images. We created a binary mask of nuclei from the MIP by normalizing the MIP intensities from 0 to 1 and applying Otsu's method of binarization with adaptive thresholding. We then filtered this mask for objects of the appropriate area and filled any voids in the mask. To identify the bright foci within nuclei, we created a separate mask from the normalized MIP using an extended maxima transform followed by a filter for objects in the binary mask of the appropriate area. Our software automatically tabulated the number of bright foci within each nucleus for each cell in the image and aggregated data for images within each well.

*G protein activation assay*

For immunoprecipitation of active Gαi3, freshly prepared cell lysates (2-4 mg) were incubated for 30 min at 4°C with the conformational Gαi3:GTP mouse antibody (1 μg) (*81*) or pre-immune control mouse IgG. Protein G Sepharose beads (GE Healthcare) were added and incubated at 4°C for additional 30 min (total duration of assay is 1 h). Beads were immediately washed 3 times using 1 ml of lysis buffer (composition exactly as above; no nucleotides added) and immune complexes were eluted by boiling in SDS as previously described. Previous work from our lab validated the use of this antibody to selectively immunoprecipitate His-Gαi3 recombinant proteins loaded with GTP (physiologic active conformation that is transient) and GTPϒS (non-hydrolysable nucleotide mimicking a stable active conformation) but not GDP (inactive conformation) (*82*).

*Homology modeling*

The prediction the protein and GIV peptide docked interface was performed by using CABSDOCK (*31*) web server and the final representation was using Pymol visualization tool (*32*). For the analysis the PDB structures 1T19, 1t2V, 1N5O were taken into consideration. 10 residues of GIV "SLSVSSDFLGKD" for their secondary structure prediction was tested on PSIPRED (*33*). The visualization of PDB:1T29 and PDB:1N5O structures was done using MolSoft LLC (https://www.molsoft.com/) .

*Image Processing*

All images were processed on ImageJ software (NIH) or FLOWJO software and assembled into figure panels using Photoshop and Illustrator (Adobe Creative Cloud). All graphs were generated using GraphPad Prism.

*Statistical Analysis and Replicates*

All experiments were repeated at least three times, and results were presented either as average ± SEM. Statistical significance was assessed using one-way analysis of variance (ANOVA) including a Tukey's test for multiple comparisons. *p < 0.05, **p < 0.01, ***p < 0.001, ****p < 0.0001.

### SUPPLEMENTARY TABLES

#### Supplementary Table 1: Summary of phenotypes

| | Property | Survival | Cell Cycle Phase arrest | Necrosis | Apoptosis | HR | NHEJ | Mutated DNA |
|---|---|---|---|---|---|---|---|---|
| **GIV +/+** | GIV can scaffold G protein and BRCA1 | + | S | - | - | - | + | - |
| **GIV -/-** | G protein and BRCA1 disconnected | - | G2/M | + | + | + | - | + |
| **GIV- WT** | GIV can scaffold G protein and BRCA1 | + | S | - | - | - | + | - |
| **GIV-F1719** | GIV binds and activates G proteins but does not bind BRCA1. | - | G2/M | + | + | + | - | + |
| **GIV-F1685** | GIV binds BRCA1 but cannot bind/activate G proteins. | - | S and G2M | + | + | + | - | + |

#### Supplementary Table 2: Summary of Predicted Nuclear Import and Export Sequences in GIV/Girdin

| Elm Name | Instances (Matched sequence) | Positions | Elm Description | Cell Compartment | Consensus Pattern | P value |
|---|---|---|---|---|---|---|
| TRG_NES_CRM1_1 | QLKAKLHD MEMERD<br>ELLQKKITN LKITCE<br>QQLESELQ DLEME<br>QLESELQD LEME<br>QLEDLEKM LKVEQE<br>ENLLDEVM KSLSVSSD | 386-399 [A]<br>642-656 [A]<br>780-792 [A]<br>781-792 [A]<br>1211-1224 [A]<br>1704-1719 [A] | Some proteins re-exported from the nucleus contain a Leucine-rich nuclear export signal (NES) binding to the CRM1 exportin protein. | nucleus, cytosol | ([DEQ].{0,1}[LIM].{2,3}[LIVMF][^P]{2,3}[LMVF].[LMIV].{0,3}[DE])\|([DE].{0,1}[LIM].{2,3}[LIVMF][^P]{2,3}[LMVF].[LMIV].{0,3}[DEQ]) | 7.626e-04 |
| TRG_NLS_MonoExtC_3 | NRKL KK | 672-677 [A] | Monopartite variant of the classical basically charged NLS. C-extended version. | nucleus, Nuclear pore, NLS-dependent protein nuclear import complex | [^DE]((K[RK])\|(RK))(([^DE][KR])\|([KR][^DE]))(([PKR])\|([^DE][DE])) | 7.252e-04 |
| TRG_NLS_MonoExtN_4 | RENRKL KK<br>KENKRL RQ | 670-677 [A]<br>836-843 [A] | Monopartite variant of the classical basically charged NLS. N-extended version. | nucleus, Nuclear pore, NLS-dependent protein nuclear import complex | (([PKR].{0,1}[^DE])\|([PKR]))((K[RK])\|(RK))(([^DE][KR])\|([KR][^DE]))[^DE] | 1.276e-03 |

### SUPPLEMENTARY FIGURES AND LEGENDS

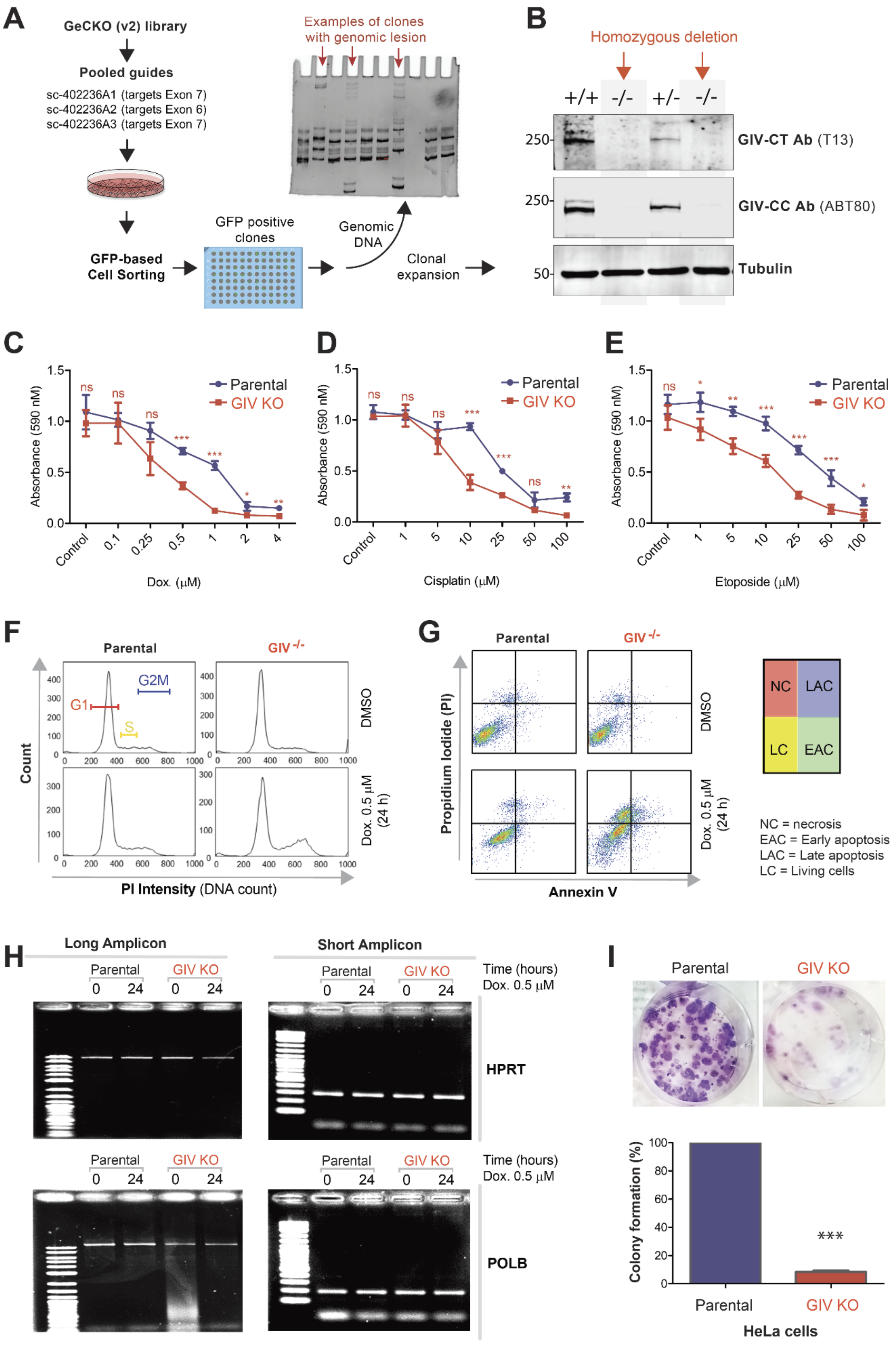


**Figure S1 [related to Figure 2]. DNA damage repair response is impaired in cells without GIV.**
**A-B.** Steps leading to creation of GIV KO HeLa lines. Cas9 target guides against GIV (CCDC88A) exons are listed in A (top). Gel showing screening of colonies by PCR of region flanking target site using genomic DNA from various cell clones from Cas9 selection (*bottom*). Immunoblots in B show examples of colonies that were either homozygous (-/-) or heterozygous (-/+) KO for GIV allele. KO clones were

pooled to avoid clonal bias. **C-E**. Line graphs display the metabolically active parental (blue) and GIV KO (red) cells that survived various doses of Doxorubicin (C), Cisplatin (D) or Etoposide (E), as determined by *MTT* tetrazolium assay (see *Methods*). Data displayed as mean ± S.E.M. and t-test was used to determine significance. (*; $p \leq 0.05$; **$p < 0.01$, ***$p < 0.001$. ns = not significant). See also **Fig 2C** for the table of IC50 values. **F**. Histograms show the percentage of cells at various stages of cell cycle (G1, S and G2/M) after challenged with Dox or vehicle control (DMSO). See bar graphs in **Fig 2D** for quantification. **G**. Necrosis (NC), apoptotic (early, EAC; late, LAC; or combined) or living cells (LC) were quantified after challenged with either Dox or vehicle control (DMSO), as assessed by annexin V staining and flow cytometry. Color coded quadrants are labeled. **H**. Long amplicon qPCR (LA-QPCR) was used to evaluate genomic DNA SB levels in control vs. GIV KO cells. Representative full-length gels showing PCR-amplified fragments of the *HPRT* (E, top panel) and *POLB* (E, bottom panel) genes. **I**. Images (top) and Bar graphs (bottom) display anchorage-dependent growth into colonies after ~2 weeks of prolonged exposure of HeLa parental and GIV KO cells to 10 nM Doxorubicin (see *Methods*).

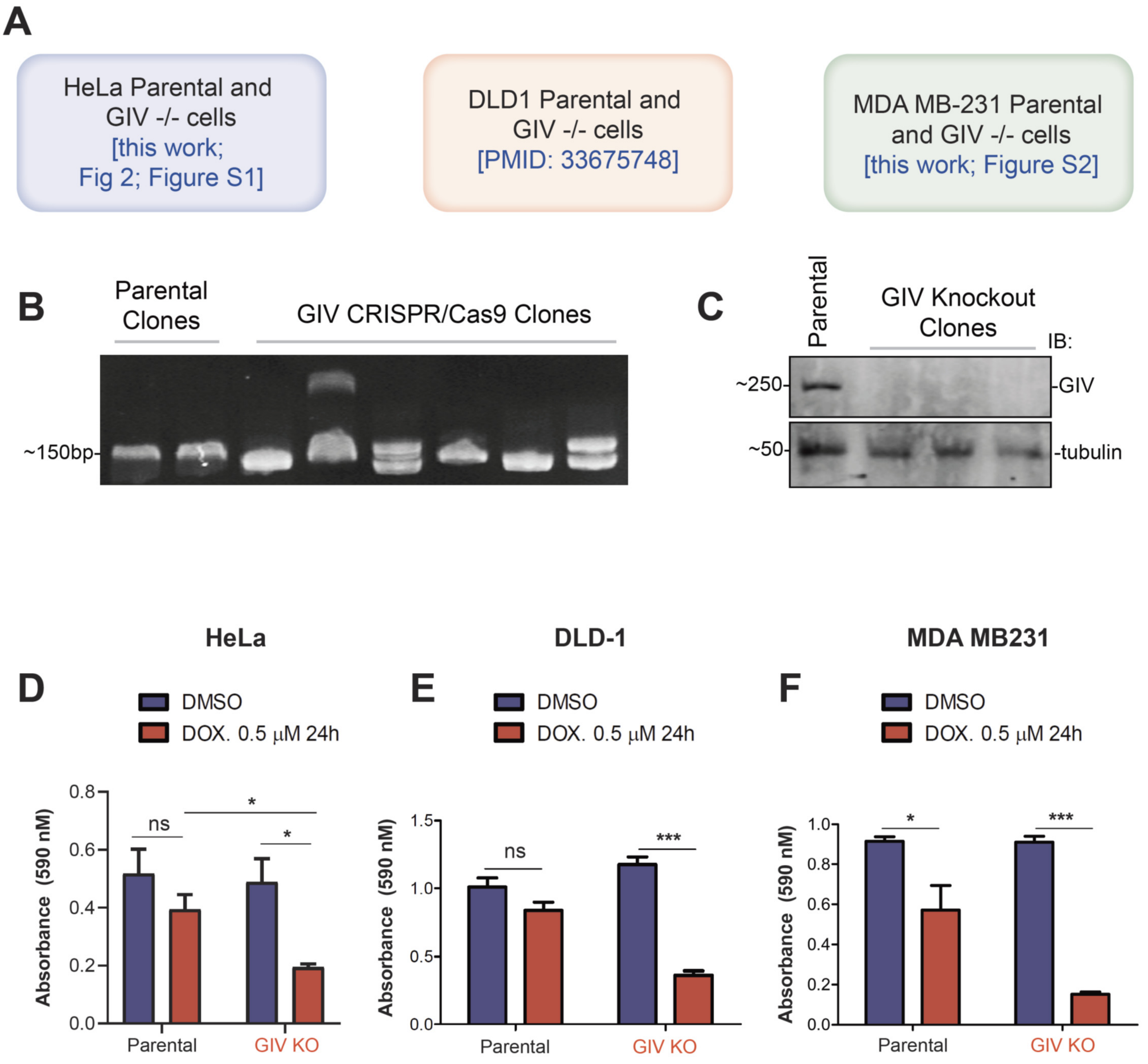


**Figure S2 [related to Figure 2]. Survival after DNA damage is impaired in DLD1 and MDA-MB-231 cells without GIV.**
**A**. Schematic outlining the three cell lines used in this work to study the role of GIV in cell survival after DNA damage. Parental and GIV KO HeLa cells are developed and validated in this work (see **Figure S1A-B**). Parental and GIV KO DLD1 cells were developed and validated in a prior work [2]. Generation and validation of parental and GIV KO MDA-MB-231 cells is described in **B-C. B-C.** Steps leading to the creation of GIV KO MDA-MB-231 lines. Cas9 target guides against GIV (CCDC88A) exons listed in **Fig S1A** (top) were used to create GIV KO lines. Gel (B) showing screening of colonies by PCR of region flanking target site using genomic DNA from various cell clones from Cas9 selection. Immunoblots in C show examples of colonies that were KO for GIV allele. KO clones were pooled to reduce clonal bias. **D-F**. Bar graphs showing % survival of metabolically active HeLa (D), DLD-1 (E) and MDA-MB-231 (F) cell lines, challenged with either Dox or vehicle control (DMSO) for 24 h, as determined by *MTT* tetrazolium *assay.* Data displayed as mean ± S.E.M. and one-way ANOVA using Tukey's multiple comparisons test was used to determine significance. (*; $p \le 0.05$; ***$p < 0.001$. ns = not significant).

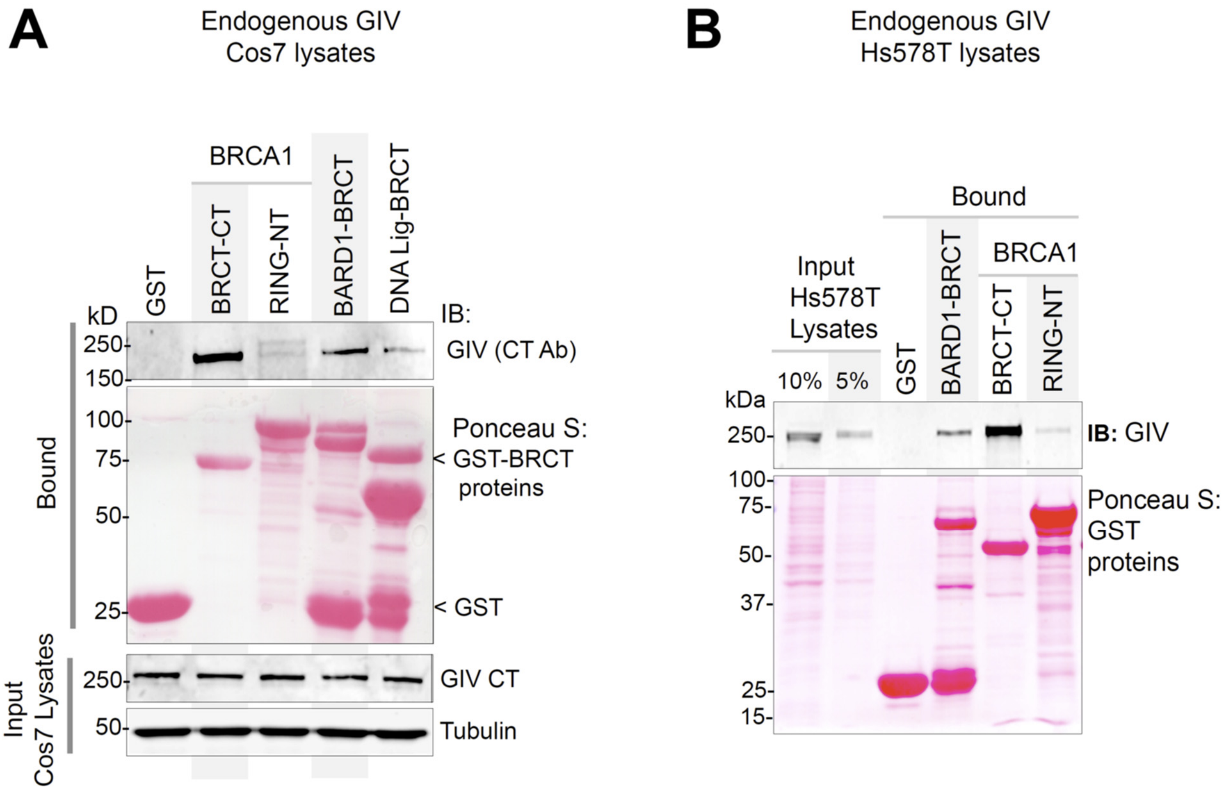


**Figure S3 [related to Figure 3]. The C-terminal BRCT module of BRCA1 binds to full length GIV from lysates of Cos7 and HeLa cells.** Pulldown assays were carried out using lysates of Cos7 (A) and Hs578T (B; a triple negative breast cancer line) cells as source of endogenous full length GIV with GST-BRCA1 and BARD1. Bound GIV was visualized by immunoblotting. See also **Fig 3D** for similar studes with lysates of HeLa cells.

**A**

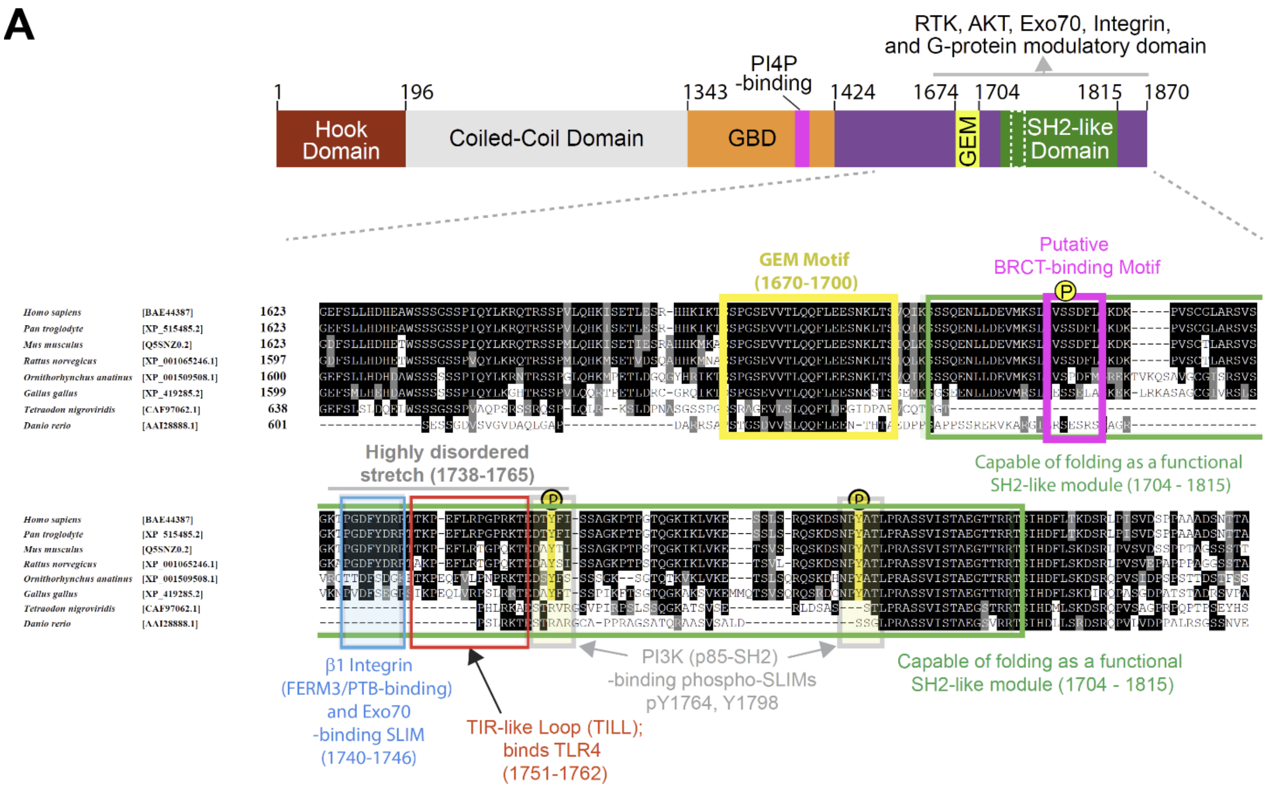


**B**

| | | | | |
|---|---|---|---|---|
| Girdin (human) | S171 6-p | VMKSLsVsSDFLGKD | Links | 1, 2, 3, 5, 7, 8, 9 |
| Girdin (mouse) | S171 6-p | VMKSLsVssDFLGKD | Links | 3, 4, 6 |

| | |
|---|---|
| 1 | Mertins P, et al. (2016) Proteogenomics connects somatic mutations to signalling in breast cancer. Nature 534 , 55 -62<br>27251275 Curated Info |
| 2 | Stuart SA, et al. (2015) A Phosphoproteomic Comparison of B -RAFV600E and MKK1/2 Inhibitors in Melanoma Cells. Mol Cell Proteomics 14 , 1599 -615<br>25850435 Curated Info |
| 3 | Mertins P, et al. (2014) Ischemia in tumors induces early and sustained phosphorylation changes in stress kinase pathways but does not affect global protein levels. Mol Cell Proteomics 13 , 1690 -704<br>24719451 Curated Info |
| 4 | Humphrey SJ, et al. (2013) Dynamic Adipocyte Phosphoproteome Reveals that Akt Directly Regulates mTORC2. Cell Metab 17 , 1009 -20<br>23684622 Curated Info |
| 5 | Zhou H, et al. (2013) Toward a comprehensive characterization of a human cancer cell phosphoproteome. J Proteome Res 12 , 260 -71<br>23186163 Curated Info |
| 6 | Wu X, et al. (2012) Investigation of receptor interacting protein (RIP3) -dependent protein phosphorylation by quantitative phosphoproteomics. Mol Cell Proteomics 11 , 1640 -51<br>22942356 Curated Info |
| 7 | Beli P, et al. (2012) Proteomic Investigations Reveal a Role for RNA Processing Factor THRAP3 in the DNA Damage Response. Mol Cell 46 , 212 -25<br>22424773 Curated Info |
| 8 | Possemato A (2009) CST Curation Set: 7403; Year: 2009; Biosample/Treatment: cell line, NCI - H2228 /untreated; Disease: non -small cell lung cancer; SILAC: -; Specificities of Antibodies Used to Purify Peptides prior to LCMS: (K/R)XX[ST]<br>Curated Info |
| 9 | Brill LM, et al. (2009) Phosphoproteomic analysis of human embryonic stem cells. Cell Stem Cell 5 , 204 - 13<br>19664994 Curated Info |

**Figure S4 [related to Figure 3]. Discovery of an evolutionarily conserved putative BRCT-binding motif on the C terminus of GIV. A.** Short linear interaction motifs (SLIMs) within GIV's C-terminus. **Top:** Bar diagram showing the various domains of GIV. GBD, G protein binding domain; GEM, Guanine nucleotide exchange modulator; SH2, Src-like homology; PI4P, phosphoinositol-4-phosphate. **Bottom**: Sequence of GIV's C-terminus showing all currently identified SLIMs. The putative BRCT-binding SLIM is highlighted in pink. **B**. A curated list of studies that reported phosphorylation at Ser1716 on GIV (Girdin). Source: Phosphosite.org, a database that was developed with grants from the NIH.

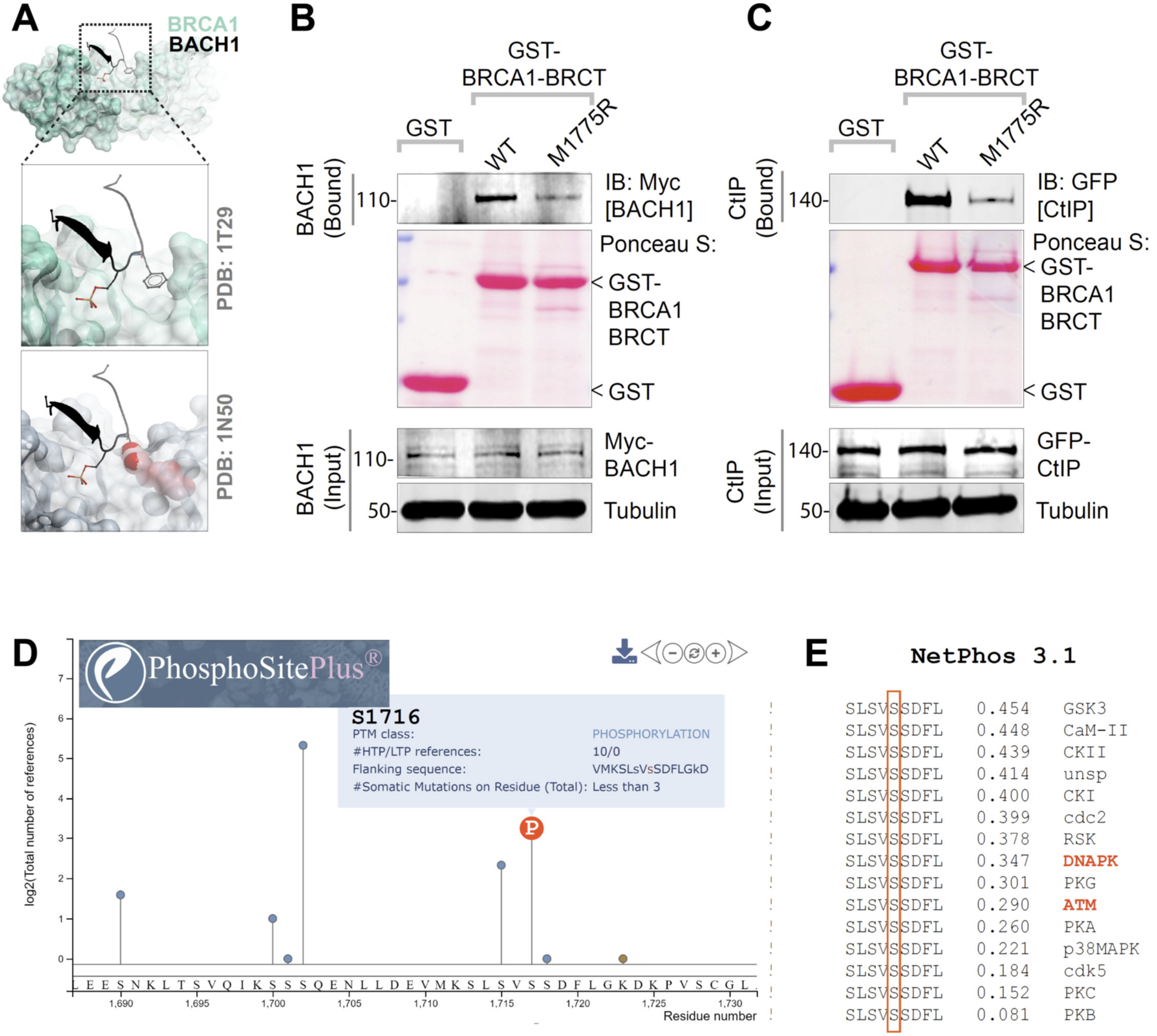


**Figure S5 [related to Figure 4]. Canonical (phosphodependent) binding of CtIP and BACH1 proteins to BRCA1 and proposed mechanism of phospho-dependent binding of GIV. A**. Top: Solved structure of BACH1-derived SxxF consensus-bearing phosphopeptide bound to BRCA1 BRCT modules. Bottom: Model of M1775R mutation in BRCA (from solved structure of the mutant BRCA1; PDB: 1N50) showing steric clash of Arg at 1775 precluding binding of Phe (F) within the SxxF motif. **B**. Pulldown assays were carried out using lysates of HEK cells as source of myc-BACH1 (B) or GFP-CtIP (C) and recombinant GST/GST-BRCA1 WT and M1775R mutant proteins. Bound proteins were visualized by immunoblotting with anti-myc (BACH1; B), or anti-GFP (CtIP; C) IgGs. **D**. A lollipop graph showing the number of independent mass spectrometry studies that reported phosphorylation at Ser1716 on GIV (Girdin). Source: Phosphosite.org, a database that was Developed with grants from the NIH. **E**. Predicted kinases that mediate such phosphorylation, as determined using the bioinformatic web-based resource, NetPhos 3.1 (https://services.healthtech.dtu.dk/).

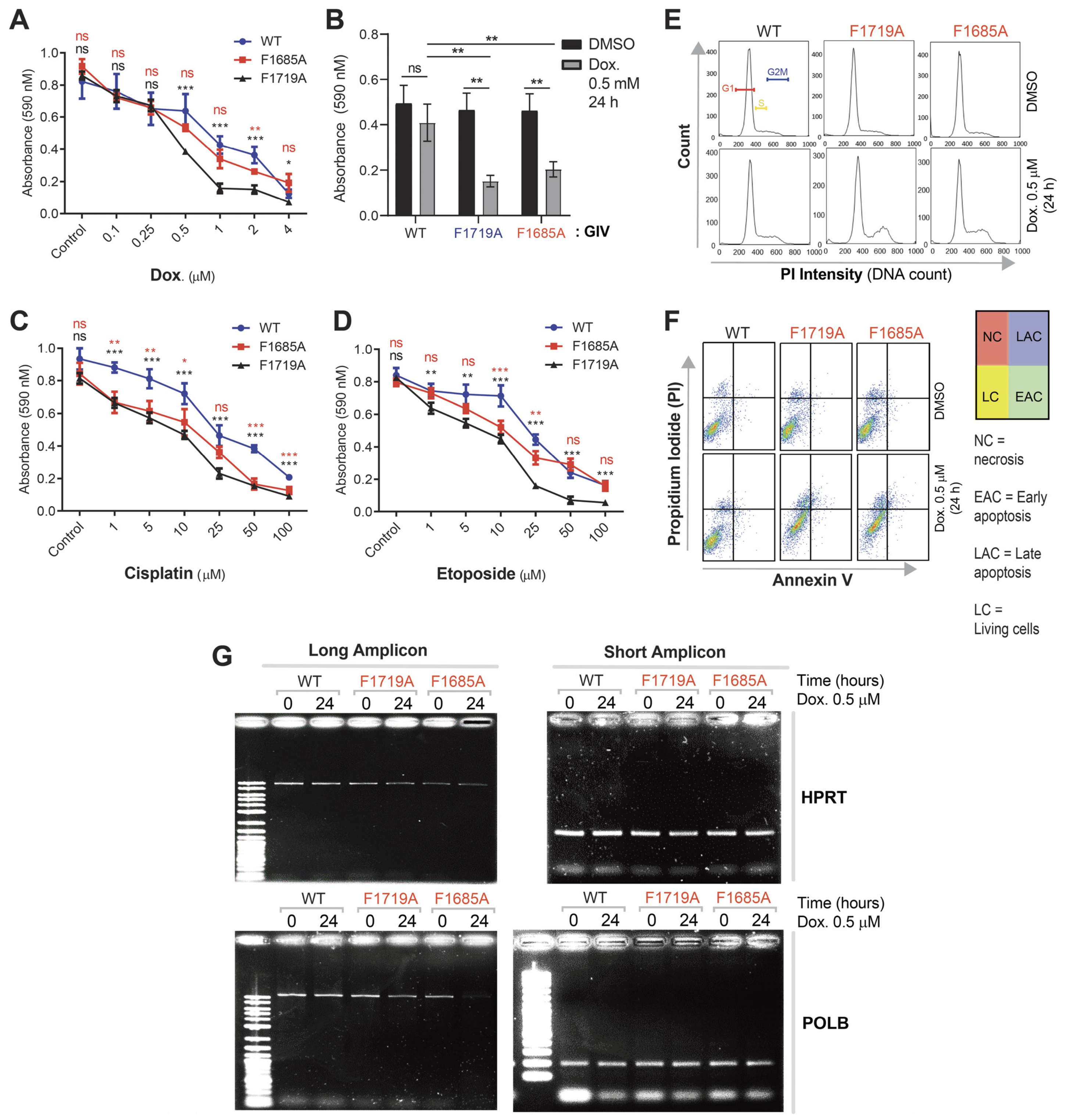


**Figure S6 [related to Figure 5]. DNA damage repair response is impaired in cells expressing mutant GIV that cannot bind BRCA1 (F1719A) or bind/activate G proteins (F1685A). A-D**. Line (A, C, D) and bar (B) graphs display metabolically active parental (blue) and GIV KO (red) cells that survived various doses of Doxorubicin (A-B), Cisplatin (C) or Etoposide (D), as determined by *MTT* tetrazolium assay (see *Methods*). Data displayed as mean ± S.E.M. and t-test was used to determine significance. (*; p ≤ 0.05; **p < 0.01, ***p < 0.001. ns = not significant). See also **Fig 5C** for the table of IC50 values. **E**. Histograms show the percentage of cells at various stages of cell cycle (G1, S and G2/M) after challenged with Dox or vehicle control (DMSO). See bar graphs in **Fig 5D** for quantification. **F**. Necrosis (NC), apoptotic (early, EAC; late, LAC; or combined) or living cells (LC) were quantified after challenged with either Dox or vehicle control (DMSO), as assessed by annexin V staining and flow cytometry. Color coded quadrants are labeled. **G**. Long amplicon qPCR (LA-QPCR) was used to evaluate genomic DNA SB levels in control vs. GIV KO cells. Representative full-length gels showing PCR-amplified fragments of the *HPRT* (E, top panel) and *POLB* (E, bottom panel) genes.

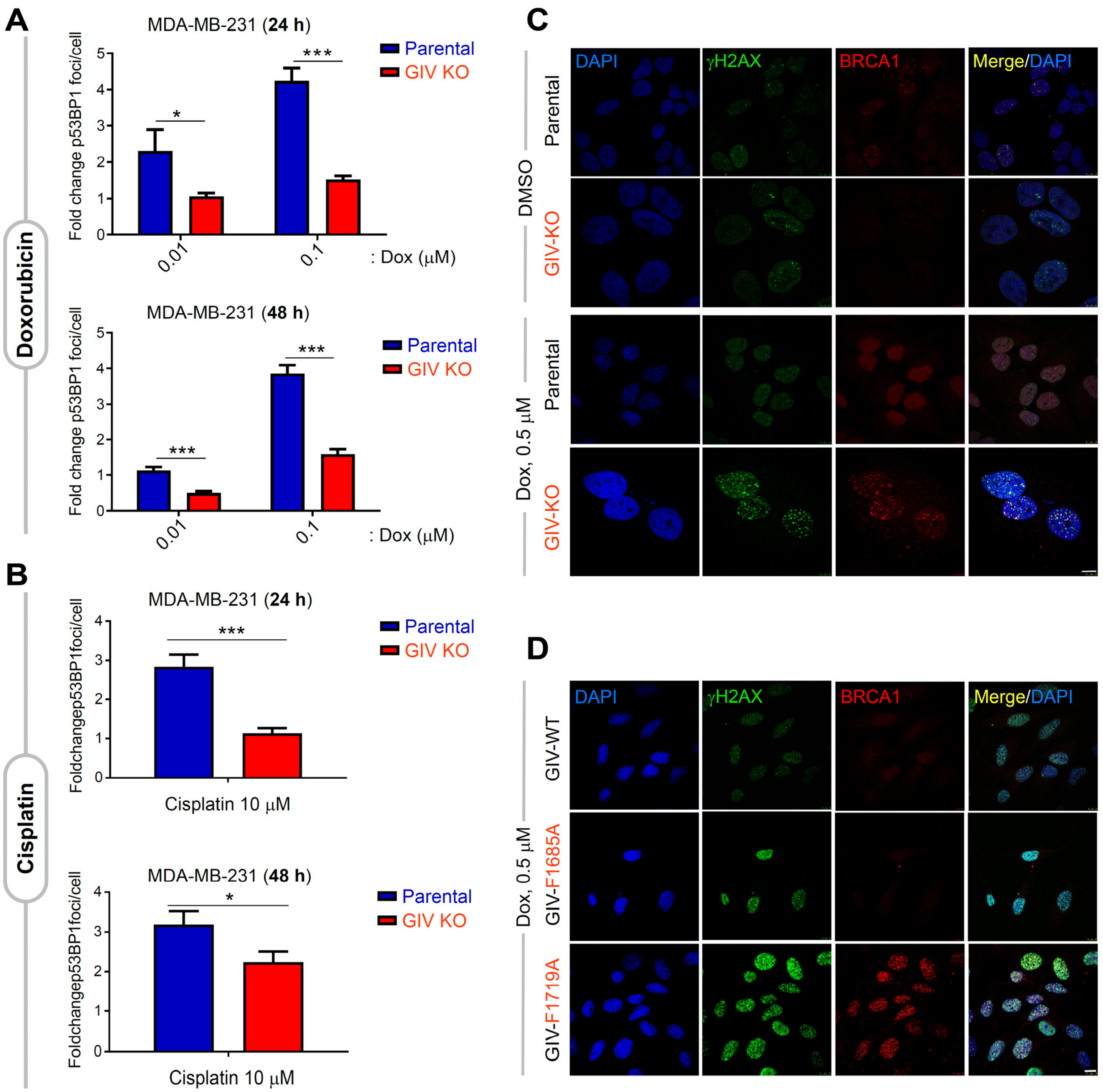


**Figure S7 [related to Figure 6]. GIV inhibits the localization of BRCA1 to sites of DNA damage. A-B.** Bar graphs display the fold change in the number of bright foci of 53BP1 in parental and GIV KO MDA-MB-231 cells stably expressing mApple-53BP1 reoprter (which detects NHEJ) upon challenge with the indicated concentrations of Doxorubicin (A) or Cisplatin (B). Data displayed as mean ± S.E.M. and t-test to determine significance. (*; $p \leq 0.05$; ***; $p \leq 0.001$). See also **Fig 6F-H** for 53BP1 reporter studies on parental and GIV KO HeLa cells. **C-D**. Control (parental) and GIV-depleted (GIV KO) HeLa cells (**C**) or GIV-depleted HeLa cells stably expressing WT or mutant GIV constructs (**D**) were challenged with Dox or vehicle control (DMSO) prior to being fixed and co-stained for ɣ2H2AX (green) and BRCA1 (red) and analyzed by confocal microscopy. Representative images are shown (scale bar = 15 µm).

### SUPPLEMENTARY EXTENDED DATA

**Extended Data 1:** Gene ontology (GO) cellular component analysis for GIV interacting proteins, as determined by DAVID GO. Data are available via ProteomeXchange with identifier PXD022601.

**Extended Data 2:** Gene ontology (GO) molecular function analysis for GIV interacting proteins, as determined by DAVID GO. Data are available via ProteomeXchange with identifier PXD02260.
